# Purine nucleosides interfere with c-di-AMP levels and act as adjuvants to re-sensitise MRSA to β-lactam antibiotics

**DOI:** 10.1101/2022.09.01.506298

**Authors:** Aaron C. Nolan, Merve S. Zeden, Christopher Campbell, Igor Kviatkovski, Lucy Urwin, Rebecca M. Corrigan, Angelika Gründling, James P. O’Gara

**Affiliations:** Microbiology, School of Biological and Chemical Sciences, University of Galway, H91 TK33, Ireland; Section of Molecular Microbiology and Medical Research Council Centre for Molecular Bacteriology and Infection, Imperial College London, London, SW7 2AZ, United Kingdom; The Florey Institute, School of Bioscience, University of Sheffield, Sheffield, S10 2TN, United Kingdom

**Keywords:** *Staphylococcus aureus*, MRSA, purine metabolism, antibiotic adjuvant, β-lactam resistance, c-di-AMP

## Abstract

Elucidating the complex mechanisms controlling *mecA*/PBP2a-mediated β-lactam resistance in methicillin resistant *Staphylococcus aureus* (MRSA) has the potential to identify new drug targets with therapeutic potential. Here, we report that mutations that interfere with *de novo* purine synthesis (*pur* operon), purine transport (NupG, PbuG and PbuX) and the nucleotide salvage pathway (DeoD2, Hpt) increased β-lactam resistance in MRSA strain JE2. Extrapolating from these findings, exogenous guanosine and xanthosine, which are fluxed through the GTP branch of purine biosynthesis were shown to significantly reduce MRSA β-lactam resistance. In contrast adenosine, which is fluxed to ATP, significantly increased oxacillin resistance, whereas inosine, which can be fluxed to ATP and GTP via hypoxanthine, only marginally reduced the oxacillin MIC. Increased oxacillin resistance of the *nupG* mutant was not significantly reversed by guanosine, indicating that NupG is required for guanosine transport, which in turn is required to reduce β-lactam resistance. Suppressor mutants resistant to oxacillin/guanosine combinations contained several purine salvage pathway mutations, including *nupG* and *hpt*. Microscopic analysis revealed that guanosine significantly increased cell size, a phenotype also associated with reduced levels of c-di-AMP. Consistent with this, guanosine significantly reduced levels of c-di-AMP, and inactivation of GdpP, the c-di-AMP phosphodiesterase negated the impact of guanosine on β-lactam susceptibility. PBP2a expression was unaffected in *nupG* or *deoD2* mutants suggesting that guanosine-induced β-lactam susceptibility may result from dysfunctional c-di-AMP-dependent osmoregulation. These data reveal the therapeutic potential of purine nucleosides as β-lactam adjuvants that interfere with the normal activation of c-di-AMP required for high-level β-lactam resistance in MRSA.

**Importance:** The clinical burden of infections caused by antimicrobial resistant (AMR) pathogens is a leading threat to public health. Maintaining the effectiveness of existing antimicrobial drugs or finding ways to reintroduce drugs to which resistance is widespread is an important part of efforts to address the AMR crisis. Predominantly the safest and most effective class of antibiotics are the β-lactams, which are no longer effective against methicillin resistant *Staphylococcus aureus* (MRSA). Here we report that the purine nucleosides guanosine and xanthosine have potent activity as adjuvants that can resensitise MRSA to oxacillin and other β-lactam antibiotics. Mechanistically, exposure of MRSA to these nucleosides significantly reduced the levels of the cyclic dinucleotide c-di-AMP, which is required for β-lactam resistance. Drugs derived from nucleotides are widely used in the treatment of cancer and viral infections highlighting the clinical potential of using purine nucleosides to restore or enhance the therapeutic effectiveness of β-lactams against MRSA and potentially other AMR pathogens.

## Introduction

*Staphylococcus aureus* is an opportunistic pathogen responsible for localised skin infections and more severe illnesses, such as bacteraemia, sepsis, infectious arthritis, pneumonia, endocarditis, urinary tract infections and toxic shock syndrome (1, 2). While significant advances have been made in our understandings of the metabolism of *S. aureus* in recent years, we have only just begun to interrogate the role of central metabolic pathways required for bacterial proliferation in the host, and their contribution to antibiotic resistance.

β-lactam antibiotics remain a gold standard for the treatment of *S. aureus* infections, while vancomycin and daptomycin are recommended for methicillin resistant *S. aureus* (MRSA) infections (3-6). However, while effective, these antibiotics are limited by slow bactericidal activity, low tissue penetration, and nephrotoxicity (7, 8). Daptomycin exhibits high protein binding and low serum concentrations which necessitates high doses of the antibiotic to achieve therapeutic levels (9). Limited treatment options for MRSA infections are exacerbated by the emergence of vancomycin intermediate *S. aureus* (VISA) strains that also display a 100-fold decrease in susceptibility to daptomycin (10, 11).

MRSA resistance to β-lactam antibiotics such as methicillin that are not cleaved by bacterial β-lactamases, is mediated through the expression of the low affinity penicillin binding protein PBP2a, encoded by the *mecA* gene located on the mobile genetic element, the staphylococcal chromosome cassette (SCC*mec*) (12, 13). The PBP2a transpeptidase crosslinks peptidoglycan in the presence of β-lactam antibiotics to confer resistance (13-15).

Carriage of SCC*mec* is linked to heterogenous, low-level resistance (HeR) to β-lactam antibiotics (13, 16). Deregulation of *mecA* transcription occurs when the MecR sensor-protease is activated by the presence of β-lactams resulting in cleavage of the MecI repressor (17). When exposed to β-lactam antibiotics, high-level, homogeneously resistant (HoR) MRSA mutants carrying accessory mutations outside SCC*mec* can also be selected (18, 19). Commonly, these mutations lead to activation of the stringent response (20, 21), increased cyclic di-adenosine monophosphate (c-di-AMP) levels (22-24), changes in the activity of RNA polymerase (6), as well as the ClpXP chaperone-protease complex (25, 26).

Nucleotide second messengers play an important role in the control of β-lactam resistance. Nucleotides are essential precursors used by all cells for DNA, RNA and signalling molecule synthesis, while the energy carriers, adenosine-triphosphate (ATP) and guanosine-triphosphate (GTP), are produced from their corresponding nucleotides adenine and guanine (27, 28). Bacteria utilise an array of nucleotide secondary messengers to regulate cellular responses to fluctuations in external stimuli and environmental conditions (29). Bacterial nucleotide messengers are typically produced from the building blocks ATP and GTP (30). Refined signal transduction pathways are necessary for *S. aureus* to survive and multiply during infection. Signal transduction in *S. aureus* influences virulence gene expression during infection, stress and antibiotic survival responses (31, 32).

*S. aureus* produces ATP/GTP through a *de novo* ten-step biosynthetic process. The *pur* operon encodes ten enzymes that first convert ribose-5-phosphate (Ribose-5-P) to inosine monophosphate (IMP); the branchpoint between ATP and GTP synthesis (Fig. 1). IMP can then be converted to adenosine monophosphate (AMP) or xanthosine monophosphate (XMP) through *purA/purB* or *guaA* respectively. AMP is then converted to adenosine diphosphate (ADP), ATP and c-di-AMP. On the alternate branch XMP is converted to guanosine diphosphate (GDP), GTP and guanosine tetraphosphate/pentaphosphate (ppGpp/(p)ppGpp) which promotes accumulation of c-di-AMP by repressing the activity of the c-di-AMP phosphodiesterase GdpP (33). Alternatively, XMP, GMP and AMP can be produced from nucleosides/nucleotides through the purine salvage pathway, including transport by the purine transporters PbuX, PbuG and NupG. There is ongoing interest in purine metabolism due to its multifaceted roles in biofilm formation, virulence and antibiotic resistance (34-36).

**Fig. 1.**
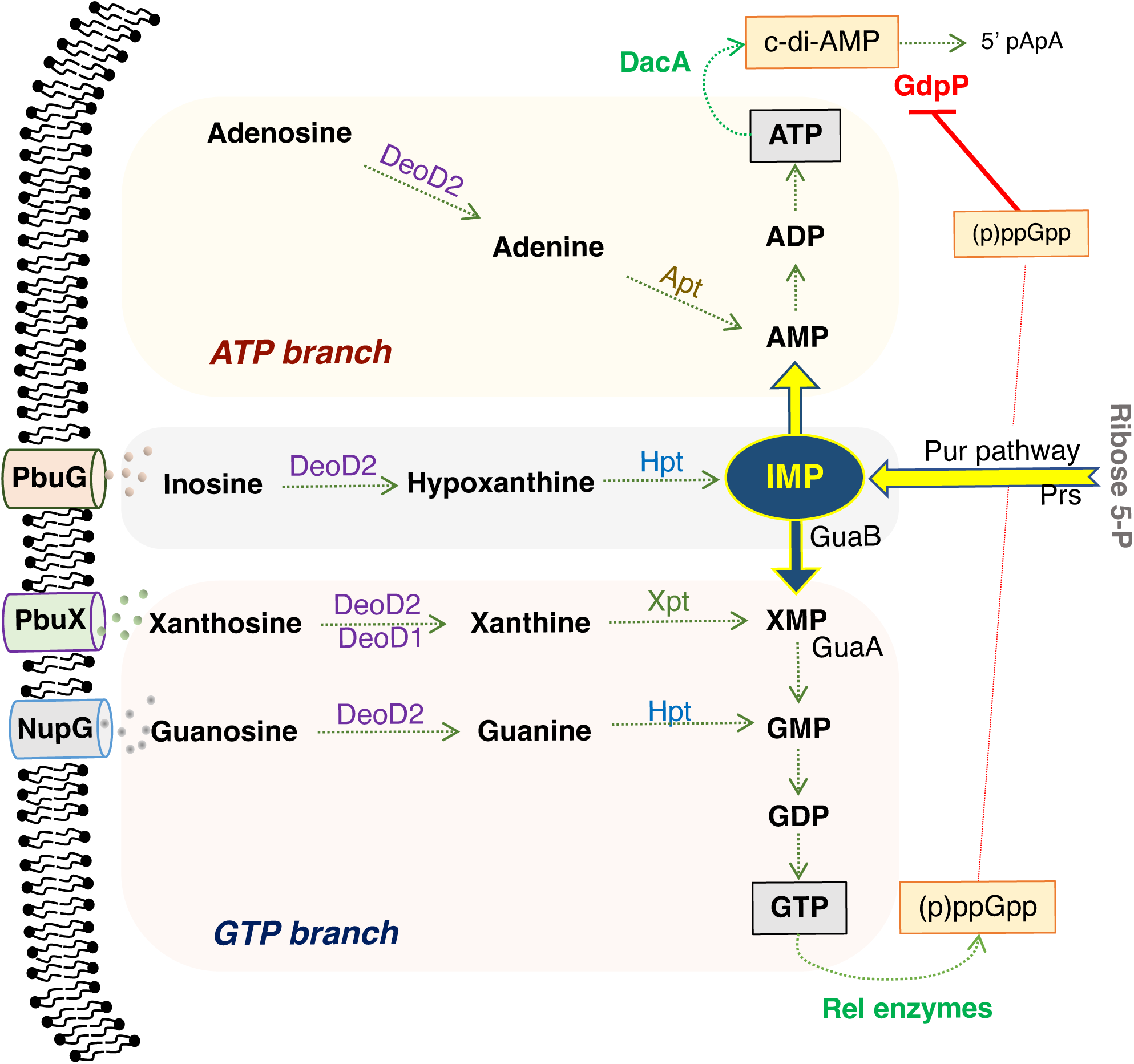
Overview of purine salvage and *de novo* biosynthetic pathways including transporters and enzymes implicated in susceptibility to nucleoside/β-lactam combinations. Inosine monophosphate (IMP) synthesized from ribose 5-P by enzymes encoded by the *pur* operon can be fluxed to ATP and GTP. Nucleotides and nucleosides can be transported into the cell via permeases including NupG, PbuX and PbuG (StgP), where they can be salvaged and returned to the intracellular nucleotide pool via the activity of enzymes including the nucleoside phosphorylases DeoD2 (SAUSA300_2091) and DeoD1 (SAUSA300_0138), hypoxanthine phosphoribosyltransferase Hpt, xanthosine phosphoribosyltransferase Xpt and adenosine phosphoribosyltransferase Apt. Mutations in *hpt, prs, guaB and guaA* have previously been implicated in the high-level, homogeneous (HoR) β-lactam resistance in MRSA (Dordel et al., 2014). GTP is a substrate for RelA, RelP and RelQ catalysed synthesis of the stringent response alarmone (p)ppGpp. The cyclic dinucleotide signalling molecule c-di-AMP is synthesized from ATP by DacA, and broken down by GdpP, whose activity is repressed by ppGpp.

The alarmones ppGpp and pppGpp play a central role in the stringent response, while c-di-AMP contributes to the regulation of cell envelope homeostasis, osmotic regulation and virulence (37-40). Altering c-di-AMP levels and/or (p)ppGpp levels lead to differences in β-lactam susceptibility of MRSA strains (31, 41). Mutation of the diadenylate cyclase gene *dacA*, leading to reduced c-di-AMP levels, resulted in the conversion of an HoR MRSA to a HeR strain (31). Conversely inactivating mutations in the c-di-AMP phosphodiesterase gene *gdpP* leading to increased c-di-AMP levels were accompanied by homogeneous resistance to methicillin (22, 23). GdpP activity is repressed by ppGpp, which is synthesised by the Rel, RelP and RelQ enzymes (41-43). In several studies it has been reported that activation of the stringent response and constitutive (p)ppGpp production, which will increase c-di-AMP levels (33), are accompanied by homogeneous methicillin resistance (21, 44). The importance of these signalling molecules to bacterial survival under different growth conditions and their absence in humans marks them as attractive antimicrobial drug targets (45).

Further understanding of molecular mechanisms to antibiotic resistance is an important part of efforts to identify new targets with the potential to re-sensitise bacteria to antibiotics including β-lactams (46). The knowledge gained from understanding such molecular mechanisms could be used to identify compounds that provide synergistic activity with β-lactams, thus providing a potential solution to antibiotic resistance and dwindling antibiotic therapy options (47-49).

In this study, the impact of exogenous purines on growth and susceptibility to β-lactams and other antibiotics is described in several strains of *S. aureus* and MRSA, and *S. epidermidis*. The contribution of several genes involved in *de novo* purine synthesis, purine transport, purine salvage, and (p)ppGpp/c-di-AMP signalling to nucleoside/β-lactam susceptibility has been elucidated. Our data reveal that re-sensitisation of MRSA to β-lactams using guanosine or xanthosine is dependent on several purine transporters and salvage pathway enzymes, and accompanied by increased cell size and reduced levels of c-di-AMP. The therapeutic potential of using purine nucleosides as β-lactam antibiotic adjuvants is discussed.

## Results

### Exogenous purines regulate MRSA growth and susceptibility to β-lactam antibiotics

The correlation between increased levels of the purine signalling nucleotides (p)ppGpp and c-di-AMP and increased resistance to β-lactam antibiotics raises the possibility that manipulation of purine metabolism (Fig. 1) by controlling the availability of purines in the growth environment can regulate the susceptibility of MRSA to β-lactam antibiotics. Exogenous purines transported into the cell are metabolised via the purine salvage pathway (Fig. 1). Growth of wild-type MRSA strain JE2 at oxacillin (OX) concentrations of 1, 16, 32 or 64 μg/ml was measured in chemically defined media without glucose (CDM) supplemented with both adenine and guanine, without adenine and guanine, with adenine alone or with guanine alone. Weak growth at the lowest OX concentration in CDM with both adenine and guanine was only measured after >30 h (Fig. S1A). This was significantly improved in CDM lacking both guanine and adenine (Fig. S1B) and growth in OX 1 μg/ml was also measured in CDM with adenine alone (Fig. S1C). Strikingly, JE2 was completely susceptible to OX in CDM supplemented with guanine alone, with no growth detected even after 48 h (Fig. S1D).

Switching to nucleosides, which are more readily solubilised in water than nucleotides, and Mueller Hinton 2% NaCl broth (MHB), which is routinely used for antibiotic susceptibility testing, addition of guanosine or xanthosine rendered JE2 susceptible to OX with the minimum inhibitory concentration (MIC) reduced from 64 to ⪕4 μg/ml (Table 1). In contrast, JE2 was significantly more resistant to OX in MHB supplemented with adenosine (MIC = 256 μg/ml, Table 1). Interestingly, supplementation of MHB with up to 1.0 g/l inosine, which can be fluxed to both the ATP and GTP branches of purine biosynthesis via hypoxanthine and IMP (Fig. 1) only marginally reduced the JE2 OX MIC to 32 μg/ml. Similarly, OX susceptibility was unchanged in MHB supplemented with both guanosine and adenosine (MIC = 64 μg/ml). Disk diffusion assays on MHA showed that exogenous guanosine also increased the susceptibility of JE2 to other β-lactam antibiotics, namely cefotaxime, cefoxitin, cefaclor and penicillin G (Fig. 2A, B), but did not show any synergy with tetracycline, vancomycin, chloramphenicol, and gentamicin (Fig. 2C).

**Table 1.**
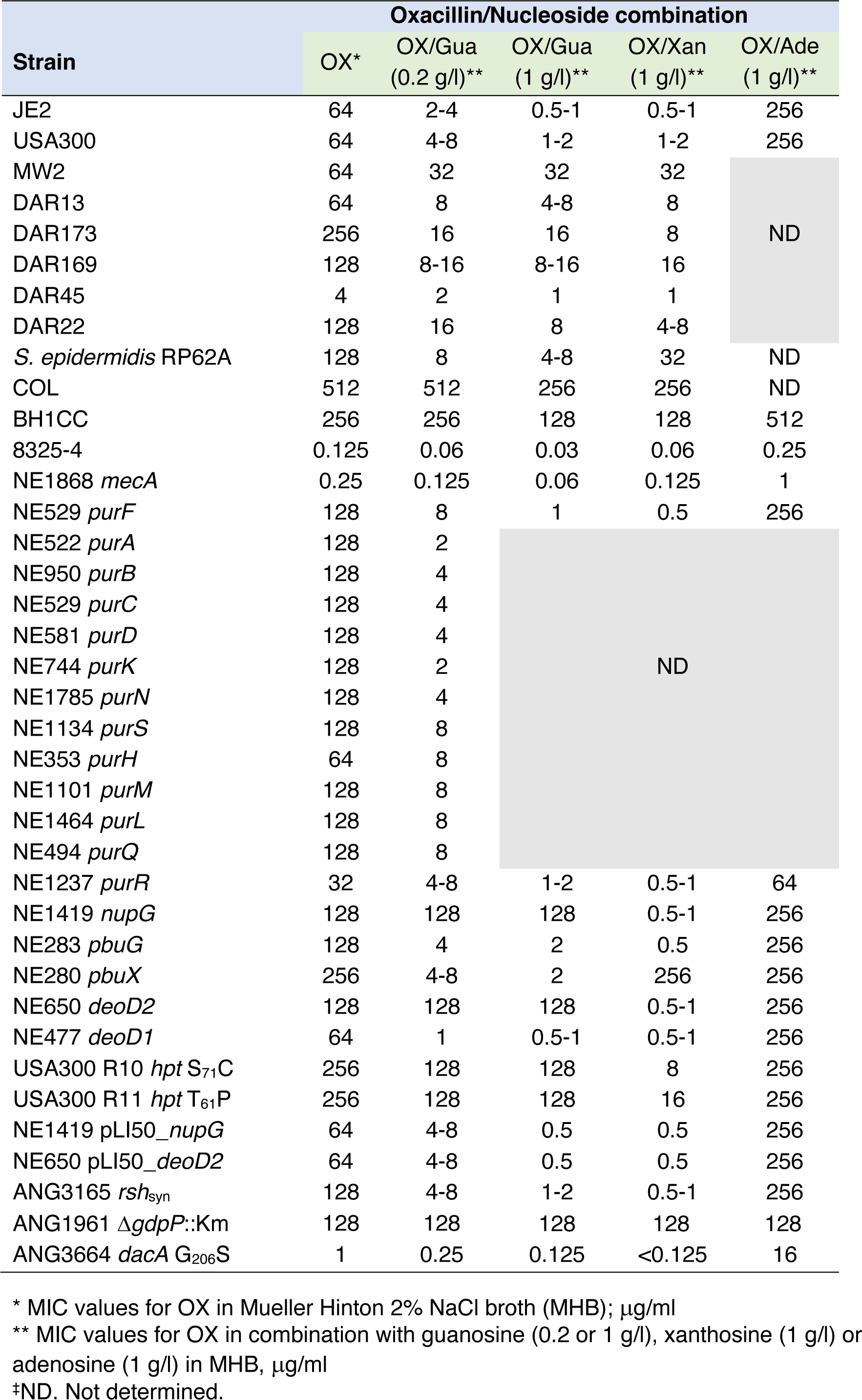
Antibacterial activity (minimum inhibitory concentrations, MICs) of oxacillin (OX) alone and in combination with guanosine (Gua), xanthosine (Xan) or adenosine (Ade) against *S. aureus* strains, purine metabolism mutants, and *S. epidermidis* RP62A.

**Fig. 2.**
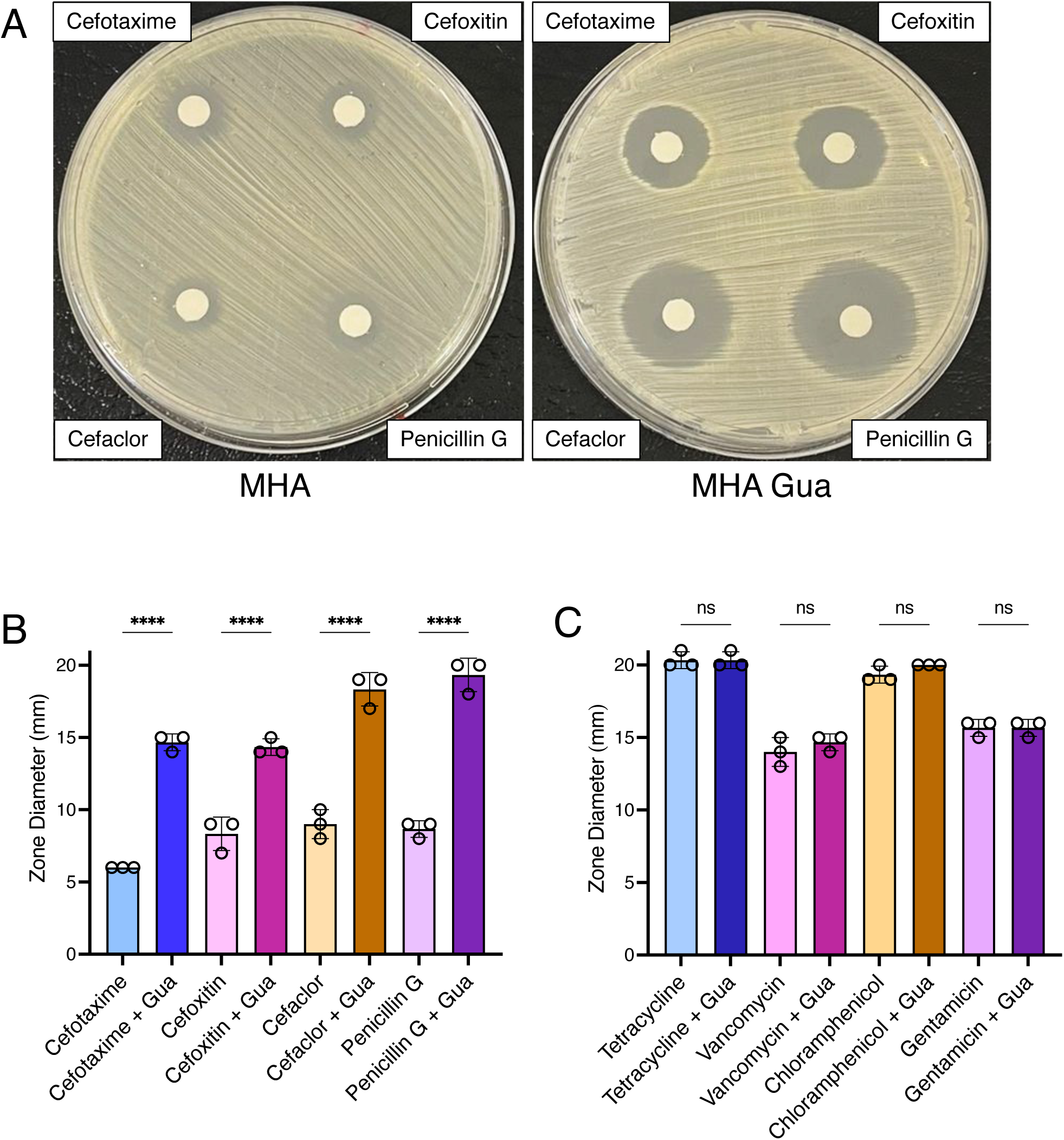
Exogenous guanosine only increases susceptibility of MRSA to β-lactams and not other classes of antibiotic. **A**. Susceptibility of wild-type JE2 to the β-lactam antibiotics cefotaxime (top left), cefoxitin (top right), cefaclor (bottom left) and penicillin G (bottom right) determined by disk diffusion assay on Mueller Hinton 2% NaCl agar (MHA) (left), or MHA supplemented with 0.2 g/l guanosine (MHA Gua) (right). This experiment was repeated three times and a representative image is shown. **B**. Susceptibility of wild-type JE2 to the β-lactam antibiotics cefotaxime, cefoxitin, cefaclor and penicillin G determined by disk diffusion assay on MHA or MHA with 0.2 g/l guanosine (Gua). **C**. Susceptibility of wild-type JE2 to tetracycline, vancomycin, chloramphenicol or gentamicin determined by disk diffusion assay on MHA, or MHA with 0.2 g/l Gua. Data (zone diameters, mm) are the average of 3 independent experiments and error bars represent standard deviation. Statistical significance was determined by one-way ANOVA. **** p<0.0001, ns, not significant

Checkerboard titration assays further revealed that guanosine increased JE2 susceptibility to OX in a concentration-dependent manner (Fig. S2). In time kill assays, combinations of guanosine and OX achieved a significant ≥2 log^10^ reduction in the number of colony forming units (CFU)/ml over 12 hours compared to OX alone, and achieved bactericidal activity at OX concentrations > 64 μg/ml (Fig. S3). At lower OX concentrations, recovery of JE2 growth after 12 hours, presumably reflected the selection and expansion of suppressors or OX HoR mutants as described previously (22, 50, 51).

This evident synergy between OX and the purine nucleosides guanosine and xanthosine versus the antagonism with adenosine and the relatively neutral effect of inosine implicate purine nucleotide homeostasis in the control of MRSA β-lactam susceptibility.

### Purine nucleoside-regulated β-lactam resistance in other *S. aureus* strains and *S. epidermidis* RP62A

Extending these analyses to other staphylococcal strains revealed that guanosine and xanthosine also reduced the OX MICs of several heterogeneously resistant MRSA strains to varying levels; USA300 (SCC*mec* type IV, CC8), MW2 (SCC*mec* type IV; CC1), DAR13 (SCC*mec* type IV; CC8), DAR173 (SCC*mec* type IV; CC5), DAR169 (SCCmec type I; CC8), DAR45 (SCC*mec* type II; CC30), DAR22 (SCC*mec* type III; CC5) and methicillin resistant *S. epidermidis* RP62A (Table 1, Table S1). Guanosine at 1 g/l was only associated with a 1-fold reduction in the OX MICs of the homogeneously resistant MRSA strains COL (SCC*mec* I, CC8) and BH1CC (SCC*mec* II, CC8) (Table 1), suggesting that genetic changes associated with high-level methicillin resistance can potentially negate the ability of purine nucleosides to reduce β-lactam resistance of MRSA. Interestingly guanosine and xanthosine also significantly reduced the OX MIC for the methicillin susceptible laboratory strain 8325-4 and the Nebraska transposon library (NTML) (52) *mecA* mutant NE1868, while adenosine increased their OX MICs indicating that therapeutic potential of purine nucleosides to control MRSA and MRSE β-lactam susceptibility is both *mecA*-dependent and *mecA*-independent.

### Mutations in the *de novo* purine biosynthesis or purine salvage pathways increase resistance to β-lactam antibiotics

Next the impacts of mutations in the *de novo* purine synthesis *pur* operon and purine salvage pathways (Fig. 1) on β-lactam resistance were investigated. NTML mutations in all tested *pur* operon genes (except *purH*) increased OX resistance, whereas mutation of the *purR* repressor gene was accompanied by increased OX susceptibility (Table 1). OX resistance was also increased in mutants of the putative transporters *nupG* (NE1419, guanine/guanosine), *pbuG/stgP* (NE283, guanine/hypoxanthine), *pbuX* (NE280, xanthine/xanthosine) and a predicted purine nucleoside phosphorylase *deoD2* (NE650) (Table 1). Consistent with the predicted roles of these nucleoside transporters, xanthosine reduced the oxacillin MIC of the *nupG* mutant and guanosine reduced OX resistance of the *pbuX* mutant (Table 1). These data suggest that NupG is the major guanosine transporter and PbuX is the main xanthosine transporter under the conditions tested. Both guanosine and xanthosine reduced OX resistance of the *pbuG* mutant (Table 1), which was previously shown to transport guanine (53). The very subtle effect of inosine on the OX MIC precluded investigation of the possible roles of NupG, PbuG and PbuX in inosine transport using susceptibility testing. Interestingly xanthosine, but not guanosine, significantly reduced the OX MIC of the *deoD2* mutant (Table 1), indicating that xanthosine can be phosphorylated by a different enzyme during purine salvage. The most likely candidate is a second putative nucleoside phosphorylase DeoD1 (NE477), which shares 67.4% identity with DeoD2 (Fig. S4A). Mutation of *deoD1* alone did not impact oxacillin susceptibility and both guanosine and xanthosine reduced the OX MIC of this mutant (Table 1), suggesting that DeoD1 and DeoD2 may be able to substitute for each other in the phosphorylation of xanthosine under specific growth conditions. Finally, in keeping with the prediction that guanine and xanthine are processed by different phosphoribosyltransferase enzymes (Hpt and Xpt, respectively, Fig. 1), xanthosine but not guanosine also reduced the OX MIC by 4-5 fold in the *hpt* mutants R10 and R11 carrying S71C and T_61_P substitutions, respectively (53) (Table 1).

To verify that the *nupG* and *deoD2* mutations were responsible for increased OX resistance in NE1419 and NE650, the *nupG*::Erm^r^ and *deoD2*::Erm^r^ alleles were transduced using phage 80α into wild-type JE2 and MW2 (Table S1). The OX MICs of the resulting transductants were the same as NE1419 and NE650, i.e. 128 μg/ml. The NE1419 and NE650 mutants were also complemented by the respective wild-type genes cloned into pLI50 (Fig. S4B and C, Table 1), restoring OX sensitivity and confirming the specific role of each in the resistance phenotype. Using Western blot analysis, no differences were detected in PBP2a expression levels in these mutants and their complemented derivatives compared to wild-type JE2 (Fig. S4D). Collectively these data are consistent with the role of purine biosynthesis and salvage in the regulation of MRSA β-lactam resistance, via a mechanism that is independent of PBP2a.

### Exogenous nucleosides control MRSA growth, cell size and β-lactam susceptibility in a NupG-dependent manner

Given their role as potential carbon sources, the impact of exogenous nucleosides on the growth of MRSA, which may in turn influence antibiotic resistance, was investigated. Weak JE2 growth in OX 1, 16 or 32 μg/ml (Fig. 3A) was completely inhibited by 0.2 g/l guanosine (Fig. 3B). In contrast, the increased OX resistance of the *nupG* mutant was not significantly reversed by guanosine (Fig. 3A, B), indicating again that NupG is the main guanosine transporter in *S. aureus* under the growth conditions tested. JE2 growth in OX was also inhibited by 0.2 g/l xanthosine (Fig. 3C), but was unaffected in MHB 0.2 g/l adenosine (Fig. 3D). Consistent with the likely role of PbuX in xanthosine transport, growth of the *nupG* mutant in OX was significantly inhibited by xanthosine (Fig. 3C), and like the wild-type was largely unaffected by adenosine (Fig. 3D).

**Fig. 3.**
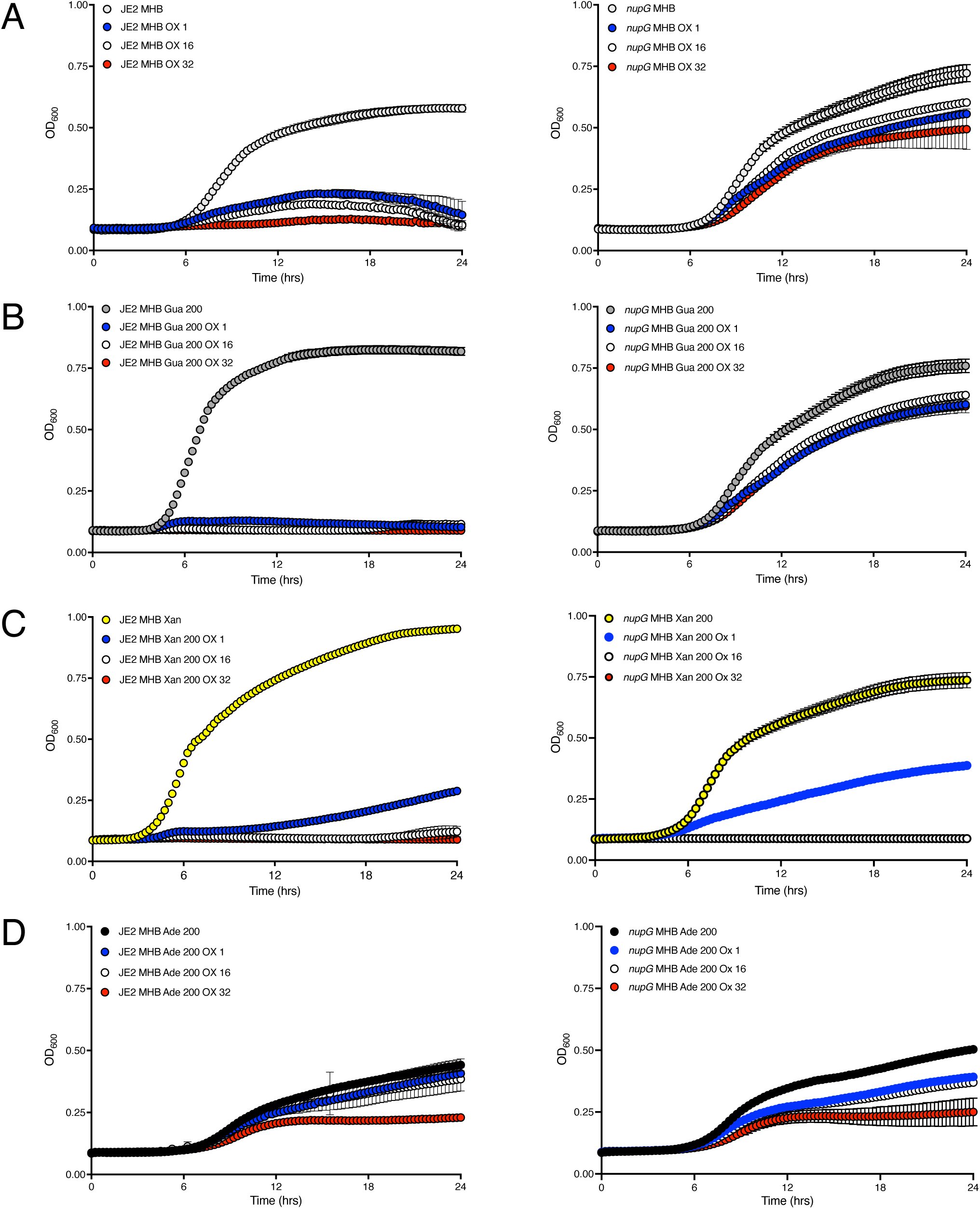
The *nupG* mutant NE1419 is more resistant to oxacillin and is not susceptible to oxacillin-guanosine combinations. **A**. Growth of wild-type and NE1419 (*nupG*) for 24 hrs at 35°C in Mueller-Hinton 2% NaCl broth (MHB) and MHB supplemented with oxacillin (OX) 1, 16 or 32 μg/ml. **B**. Growth of wild-type and NE1419 (*nupG*) in MHB Gua 0.2g/l and MHB Gua 0.2 g/l supplemented with OX 1, 16 or 32 μg/ml. **C**. Growth of wild-type and NE1419 (*nupG*) in MHB Xan 0.2 g/l and MHB Xan 0.2 g/l supplemented with OX 1, 16 or 32 μg/ml. **D**. Growth of wild-type and NE1419 (*nupG*) in MHB Ade 0.2 g/l and MHB Ade 0.2 g/l supplemented with OX 1, 16 or 32 μg/ml. Growth (OD_600_) was measured at 15 min intervals in a Tecan plate reader. Data are the average of 3 independent experiments and error bars represent standard deviation.

Strikingly, guanosine significantly increased the optical density (OD_600_) of JE2 cultures, whereas adenosine negatively impacted growth and xanthosine had no effect (Fig. 4A). However, enumeration of JE2 colony forming units (CFUs) revealed no significant differences between MHB or MHB supplemented with guanosine, xanthosine or adenosine (Fig. 4B). For the *nupG* mutant, exogenous guanosine or xanthosine did not significantly affect growth, but cell density (OD_600_) was significantly reduced by adenosine (Fig. 4C). CFU numbers for *nupG* cultures were unaffected by any nucleoside (Fig. 4D). Given that all three of these ribose-containing nucleosides have the potential to serve as carbon sources, their different effects on cell density indicates that their impact on β-lactam susceptibility was not a nutritional effect.

**Fig. 4.**
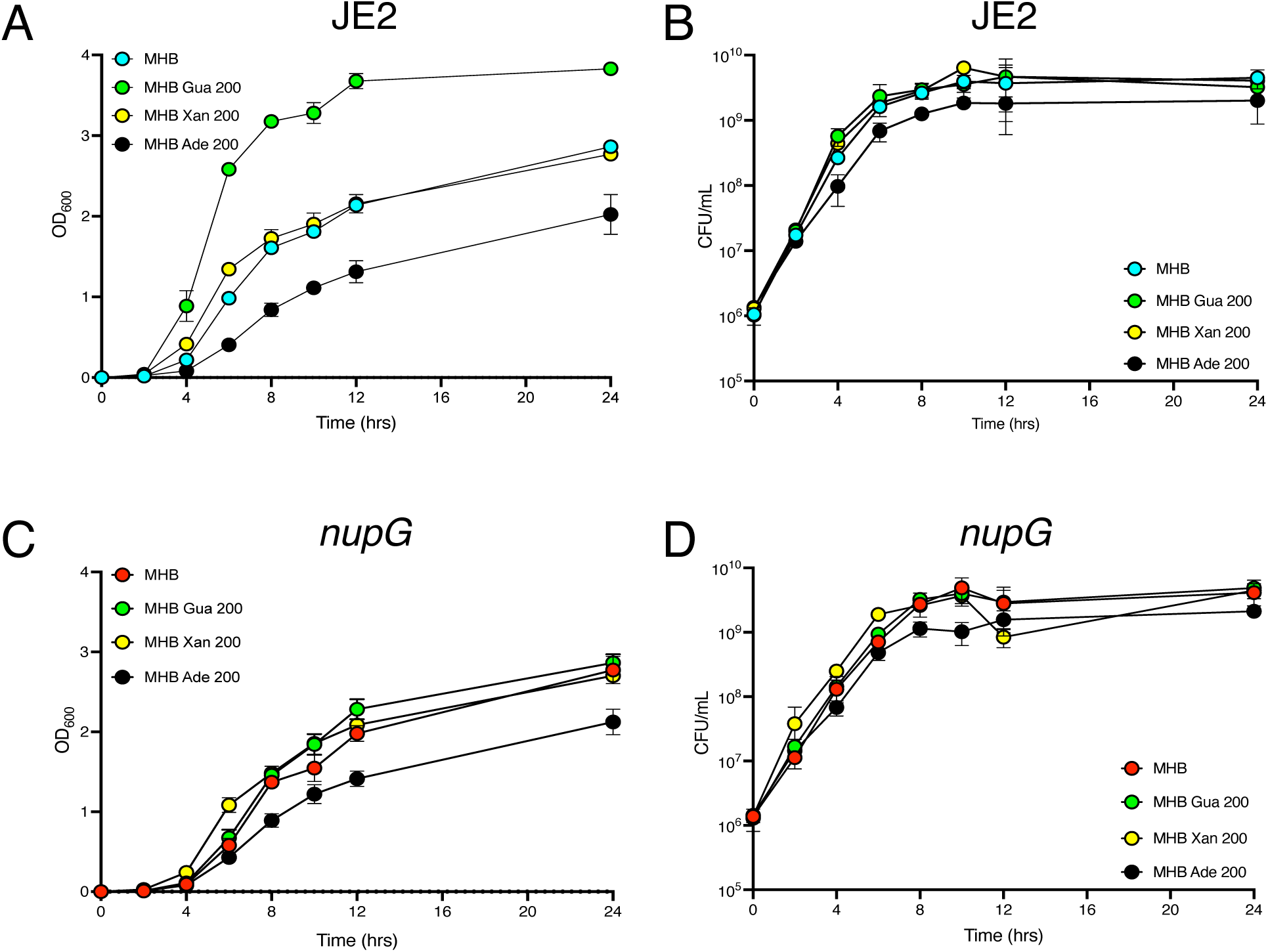
Exogenous guanosine, xanthosine and adenosine impact growth and oxacillin susceptibility in a NupG-dependent manner. Growth of wild-type and NE1419 (*nupG*) at 35°C in Mueller-Hinton 2% NaCl broth (MHB) or MHB supplemented with 0.2 g/l guanosine (Gua), Xanthosine (Xan) or adenosine (Ade) was compared by measuring cell density (OD_600_) **(A and C)** or enumerating colony forming units (CFU/ml) **(B and D)** after 0, 2, 4, 6, 8, 12 and 24 h. Both strains were grown in 25 ml culture volumes in 250 ml flasks (1:10 ratio). Data are the average of 3 independent experiments and error bars represent standard deviation.

A recent study revealed that exposure to OX increases *S. aureus* cell size and leads to cell division defects (54). To investigate the impact of exogenous guanosine on morphology and cell size, which can impact growth culture optical density, confocal microscopy experiments were performed on cells collected from JE2 cultures grown in MHB NaCl, MHB NaCl OX, MHB NaCl Gua and MHB NaCl OX/Gua and stained with Vancomycin-BODIPY or WGA Alexa Fluor 594 (Fig. 5A, B). The diameter of cells grown in guanosine was slightly, yet significantly, increased compared to cells grown in MHB NaCl only. Cell size was further increased in cells grown in oxacillin, with the largest cell size measured in the OX/Gua combination (Fig. 5A, B). Increased cell size may explain, at least in part, why exogenous guanosine increased the optical density (OD_600_) of JE2 cultures, independent of changes in the number of CFUs. Furthermore, the dramatic effect of the OX/Gua combination on both increased β-lactam susceptibility and cell size raised the possibility that these phenotypes are associated with altered regulation of c-di-AMP, which as noted earlier is known to control both of these phenotypes (37-40).

**Fig. 5.**
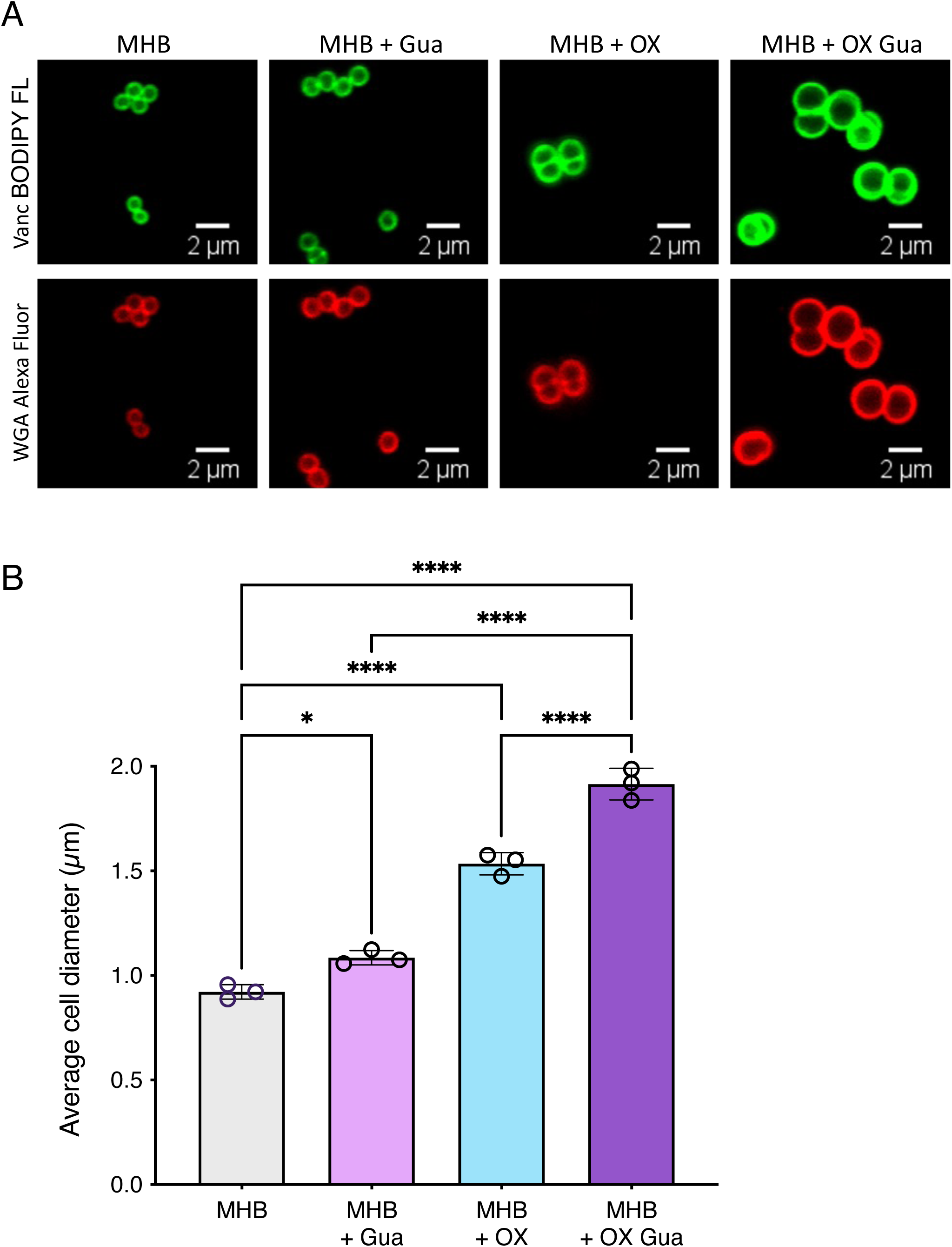
MRSA cell size is dramatically increased by growth in exogenous guanosine and oxacillin. **A**. Representative microscopic images of JE2 cells grown in MHB NaCl or MHB NaCl supplemented with guanosine (Gua, 0.2 g/l), oxacillin (OX,1 μg/ml) or a combination of Gua and OX labelled with vancomycin BODIPY FL (green, top panel) or WGA Alexa Fluor 594 (red, bottom panel). **B**. Average diameter of JE2 cells grown in MHB NaCl or MHB NaCl supplemented with Gua (0.2 g/l), OX (1 μg/ml) or a combination of Gua and OX. Images of cells from three biological replicates were acquired using Fv3000 confocal microscope and software, 40 cells measured per biological replicate (120 cells in total per condition), and the average and standard deviations for the three biological replicates were plotted using GraphPad Prism V9. Asterisks indicate statistically significant difference according to one-way ANOVA followed by Tukey’s multiple comparison post-hoc test (* p<0.05, ** p<0.01, *** p<0.001, **** p<0.0001). Error bars indicate standard deviation.

### Suppressor mutations in purine metabolism, transcriptional and translational regulation and ClpX overcome MRSA susceptibility to oxacillin/purine combinations

Repetitive passaging experiments in MHB 2% NaCl supplemented with OX (32 μg/ml) and guanosine (0.2 g/l) were used to isolate suppressor mutants resistant to the OX/Gua combination. For control purposes, HoR mutants isolated from JE2 grown in medium supplemented with OX alone (32 μg/ml) were also isolated. In total 16 OX/Gua suppressor mutants and 4 HoR mutants that exhibited stable resistance to oxacillin were characterised and sequenced (Table 2). All 16 OX/Gua suppressor mutants exhibited higher MICs to OX, OX/Gua and OX/Xan combinations (Table 2). Several mutations directly impacted purine biosynthetic and salvage pathways supporting the hypothesis that purine homeostasis controls β-lactam resistance. The identification of a W_29_Stop suppressor mutation in *nupG* (GS28) is consistent with the phenotype of the *nupG* transposon mutant (NE1419) described above. A *hpt* L_113_Stop mutation was identified in suppressors GS8 and GS25 (Table 2) (Fig. 1). Consistent with the increased β-lactam resistance of GS8 and GS25, the R10 and R11 *hpt* mutants (53) were also found to have an increased oxacillin MIC = 256 μg/ml (Table 1). However, in contrast the R10 and R11 *hpt* mutants (53) (Table 1), the oxacillin MIC of the GS8 and GS25 *hpt* null mutants was not substantially reduced by xanthosine (Table 2), presumably due to the loss of Hpt activity in G8 and GS25 versus altered Hpt activity of R10 and R11. Suppressor mutants GS15, GS16, GS22 and GS23 carry G_35_D mutations in ribose-phosphate pyrophosphokinase (Prs), which is responsible for the synthesis of phosphoribosyl diphosphate (PRPP), the substrate for the *pur* operon-encoded enzymes. Mutations that may phenocopy increased activity of the stringent response required for expression of high level β-lactam resistance include RpoB mutations (T_480_K in GS4, GS29 and T_622_I in GS10, GS17 and GS28) that lead to increased β-lactam resistance, increased doubling times and thickened cell walls similar to increased Rel activity (44, 55-57), as well as mutations that potentially impact translation (tRNA-Met in GS3, GS6, GS21, tRNA-Leu in GS10 and tRNA-Ala in GS21, GS27). GS27 has a G_278_Stop substitution in ClpX, which is known to control β-lactam resistance (25). GS28 has a A_344_T substitution in FemA previously implicated in resistance (58), as well as additional mutations in *rpoB* and *nupG*. Similarly, the PdhB G_65_V mutations in GS16 and GS23 occur in conjunction with Prs G_35_D substitutions and may further enhance growth and resistance in media supplemented with OX/Gua. Mutations in *pdhB* were previously shown to confer a growth advantage under osmotic stress (59).

**Table 2.** Antibacterial activity (minimum inhibitory concentrations, MICs) of oxacillin (OX) alone and in combination with guanosine (Gua) or xanthosine (Xan), against wild-type MRSA and selected OX/Gua and OX (HoR) suppressor mutants.

| Antibiotic →<br>↓ Strain | OX<sup>1</sup> | OX/Gua<br>(1 g/l)<sup>2</sup> | OX/Xan<br>(1 g/l)<sup>2</sup> | Mutation(s) |
| --- | --- | --- | --- | --- |
| **Wild-type** |  |  |  |  |
| JE2 | 64 | 0.5-1 | 0.5-1 | -<sup>3</sup> |
| **OX/Gua suppressors** |  |  |  |  |
| GS1 | 256 | 256 | 128 | ClpX, G<sub>278</sub>Stop |
| GS3 | 256 | 128 | 128 | tRNA-Met (RS09985) |
| GS4 | 256 | 256 | 128 | RpoB, T<sub>480</sub>K |
| GS6 | 256 | 256 | 128 | tRNA-Met (RS09985) |
| GS8 | 256 | 256 | 128 | Hpt, L<sub>113</sub>Stop |
| GS10 | 256 | 256 | 128 | RpoB, T<sub>622</sub>I;<br>tRNA-Leu (RS10005) |
| GS15 | 256 | 256 | 128 | Prs, G<sub>35</sub>D |
| GS16 | 256 | 256 | 128 | Prs, G<sub>35</sub>D;<br>PdhB G<sub>65</sub>V |
| GS17 | 256 | 128 | 128 | RpoB, T<sub>622</sub>I |
| GS21 | 256 | 128 | 128 | tRNA-Ala (RS02670);<br>tRNA-Met (RS09985) |
| GS22 | 256 | 256 | 128 | Prs, G<sub>35</sub>D;<br>RS06410, G<sub>108</sub>Stop |
| GS23 | 256 | 256 | 128 | Prs, G<sub>35</sub>D;<br>PdhB G<sub>65</sub>V |
| GS25 | 256 | 256 | 256 | Hpt, L<sub>113</sub>Stop |
| GS27 | 256 | 128 | 128 | ClpX, G<sub>278</sub>Stop;<br>tRNA-Ala (RS02670) |
| GS28 | 256 | 256 | 128 | RpoB, T<sub>622</sub>I;<br>NupG, W<sub>29</sub>Stop;<br>FemA, A<sub>344</sub>T |
| GS29 | 256 | 128 | 128 | RpoB, T<sub>480</sub>K;<br>Intergenic |
| **OX suppressors (HoRs)** |  |  |  |  |
| OS14 | 128 | 256 | 256 | ClpX, G<sub>98</sub>Stop |
| OS18 | 128 | 64 | 32 | RsbW, P<sub>101</sub>H |
| OS26 | 128 | 4 | 4 | PBP2, F<sub>44</sub>S |
| OS30 | 128 | 64 | 32 | RsbW, P<sub>101</sub>H |

<sup>1</sup> MICs for OX; µg/ml
<sup>2</sup> MICs for OX in combination with 1g/l guanosine or xanthosine; µg/ml
<sup>3</sup> A HemE L<sub>111</sub>F mutation was identified in JE2 and all of the GS and OS suppressors sequenced in this experiment.

The 4 Ox HoRs isolated for control purposes displayed varying MICs for OX/Gua and OX/Xan, although all were more resistant to the combinations than the wild-type JE2 strain (Table 2). OS14, which has ClpX G_98_Stop mutation was highly resistant to the OX/Gua combination (MIC = 256 μg/ml), and even more resistant to the OX/Xan combination (MIC = 256 μg/ml) than the GS1 and GS27 suppressor mutants with ClpX G_278_Stop mutations (MIC = 128 μg/ml). OS18 and OS30 were found to have P_101_H substitutions in RsbW, the anti-sigma factor that controls the activity of the alternative sigma factor SigB, indicating that, consistent with a previous report (60), an altered σ^B^-mediated stress response can increase β-lactam resistance and partially overcome susceptibility to the OX/Gua and OX/Xan combinations. Finally, a PBP2 F44S substitution in OS26 was associated with a modest increase in resistance to oxacillin, and the OX/Gua and OX/Xan combinations (Table 2). Overall these data reveal that aside from suppressors in the purine salvage pathway, mutations associated with the HoR phenotype can also confer resistance to OX/nucleoside combinations.

### Guanosine-oxacillin combination treatment is associated with a *nupG* and *deoD2*-dependent reduction in c-di-AMP levels

The ATP and GTP-derived stringent response alarmone (p)ppGpp and nucleotide second messenger c-di-AMP regulate β-lactam resistance in MRSA (23, 31). Comparison of GTP levels using a GTPase-Glo bioluminescence assay surprisingly revealed no significant differences between JE2, *nupG* and *deoD2* grown in MHB NaCl supplemented with guanosine (0.2 g/l) and/or oxacillin (1 or 32 μg/ml) (Fig. S5A). Using the same strains and growth conditions, radiochemical thin layer chromatography (TLC) assays also to our surprise revealed no significant differences in (p)ppGpp levels, except in JE2 grown in MHB NaCl supplemented with guanosine (Fig. S5B). The absence of significant changes in the stringent response alarmone in the *nupG* or *deoD2* mutants or during growth in OX/Gua combinations revealed that guanosine/xanthosine-mediated increase in oxacillin susceptibility is independent of (p)ppGpp signalling. Furthermore, the oxacillin MIC of a *rsh*syn mutant lacking three conserved amino acids in the (p)ppGpp synthetase domain of the RSH (Rel) enzyme (33) was decreased and increased by guanosine/xanthosine and adenosine, respectively, similar to the wild-type (Table 1) further suggesting that these phenotypes are (p)ppGpp-independent.

A competitive ELISA assay revealed significantly reduced c-di-AMP levels in the *nupG* and *deoD2* mutants grown in MHB NaCl (Fig. 6A) without oxacillin supplementation, which may be in keeping with the disruption to the purine salvage pathway cause by these mutations. Consistent with the putative role of NupG as a guanosine permease, exogenous guanosine (0.2 g/l) did not impact c-di-AMP production in the *nupG* mutant (Fig. 6A). Interestingly exogenous guanosine did restore c-di-AMP to wild-type levels in the *deoD2* mutant (Fig. 6A), suggesting that even though this was not associated with any change in oxacillin susceptibility (Table 1), the combination of NaCl and imported guanosine leads to increased c-di-AMP production. This experiment was repeated using MHB media without 2% NaCl (Fig. S6), which also showed that c-di-AMP was reduced in the *nupG* and *deoD2* mutants, and that exogenous guanosine restored c-di-AMP to wild-type levels in both mutants. Collectively, these data reveal that *nupG* and *deoD2* mutations have robust impact on c-di-AMP in MHB and in MHB NaCl. However, the data suggest that in the absence of NaCl, and oxacillin, alternative permeases and nucleoside phosphorylases may substitute for the absence of NupG and DeoD2 respectively, and restore c-di-AMP regulation and homeostasis. Furthermore, NupG is a predicted sodium-nucleoside symporter, suggesting that in the absence of sodium it will not import its substrate nucleoside.

**Fig. 6.**
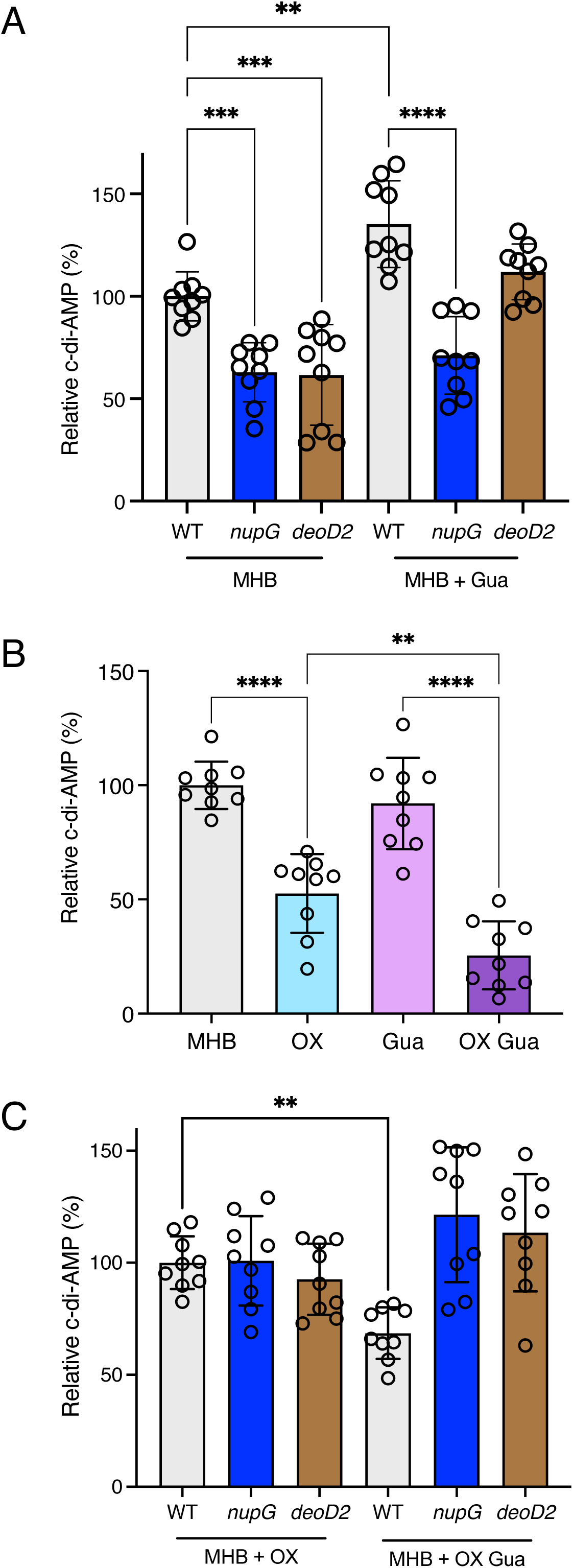
Exogenous guanosine controls c-di-AMP in a *nupG* and *deoD2*-dependent manner. **A**. Relative c-di-AMP levels in wild-type, *nupG* and *deoD2* grown for 18 h in 5 ml cultures with and without 0.2 g/l guanosine (Gua). **B**. Relative c-di-AMP levels in wild-type grown for 18 h in 12 ml cultures with and without 0.2 μg/ml oxacillin (Ox) and/or 0.2 g/l Gua. **C**. Relative c-di-AMP levels in wild-type, *nupG* and *deoD2* grown for 18 h in 12 ml cultures with and without 0.2 μg/ml Ox and/or 0.2 g/l Gua. Relative c-di-AMP levels are the averages of nine biological replicates from a combination of three separate experiments with three bio-replicates each, normalized as % relative to wild-type. Asterisks indicate statistically significant difference according to one-way ANOVA followed by Tukey’s multiple comparison post-hoc test (* p<0.05, ** p<0.01, *** p<0.001, **** p<0.0001). Error bars indicate standard deviation.

Next, the levels of c-di-AMP were measured in the wild-type, *nupG* and *deoD2* strains grown in MHB NaCl with OX, which are the same culture conditions used to measure OX susceptibility. Given the negative effect of OX/Gua combinations on growth, the culture volumes were scaled up for these experiments from 5 ml in 20 ml tubes (Fig. 6A) to 12 ml in 125 ml flasks (Fig. 6B). In contrast to growth in tubes (Fig. 6A), growth in larger cultures volumes and flasks did not reveal any increase in wild-type c-di-AMP levels in MHB NaCl supplemented with guanosine without OX (Fig. 6B), further showing that changes in growth conditions can lead to differentially regulated c-di-AMP levels in the cells. Importantly, the guanosine-induced OX susceptibility phenotype of the wild-type correlated with the lowest levels of c-di-AMP compared to growth in guanosine or OX alone (Fig. 6B). This significant effect was evident even when very low, sub-MIC concentration (0.2 μg/ml) of OX used to minimise the effect on growth. In contrast, growth of the *nupG* and *deoD2* mutants in MHB supplemented with guanosine and OX increased c-di-AMP levels, albeit not reaching statistical significance (Fig. 6C). The reduced levels of c-di-AMP in MRSA cells exposed to the OX/Gua combination observed in this experiment is consistent with the significant increase in cell size observed the same growth conditions (Fig. 5).

Consistent with an important role for c-di-AMP in these phenotypes, the increased OX resistance of a Δ*gdpP*::Km mutant (ANG 1961) in which c-di-AMP levels are increased (23) was unaffected by guanosine, xanthosine or adenosine even at 1 g/l (Table 1). Similarly, the low OX MIC of a *dacA* G_206_S mutant in which c-di-AMP synthesis is impaired but not abolished (61) was reduced further by exogenous guanosine or xanthosine, and increased significantly by adenosine (Table 1). Overall these data indicate that exogenous guanosine interferes with the normal activation of c-di-AMP production required for full expression of β-lactam resistance in MRSA.

## Discussion

Identification and characterisation of antibiotic adjuvants to maintain the therapeutic effectiveness of currently licenced antibiotics, or facilitate the reintroduction of drugs for which resistance is widespread, is a promising part of efforts to address the AMR crisis. In keeping with previous studies implicating purine metabolism in the regulation of MRSA β-lactam resistance (44, 62), we report here that mutations interfering with the purine salvage and biosynthetic pathways increased oxacillin resistance. These data prompted us to evaluate if the opposite was true and if susceptibility to β-lactams could be increased by exposing MRSA to exogenous purine nucleotides and nucleosides. Guanosine and xanthosine, which are fluxed through the GTP branch of purine biosynthesis reduced the oxacillin MIC of JE2 into the susceptibility range, in a dose dependent manner, whereas adenosine, which is fluxed to ATP, significantly increased oxacillin resistance. Furthermore inosine, which can be fluxed to ATP or GTP had almost no effect. These data reveal an important role for purine homeostasis in controlling the susceptibility of MRSA to β-lactam antibiotics and add to several recent studies implicating the regulation of purine biosynthesis in the control of virulence (35, 63, 64). Interestingly, as part of a 2017 study from our laboratory showing that β-lactams can attenuate MRSA virulence, transcriptomic analysis revealed that, after the *agr* locus, the *pur* operon was the second most down-regulated gene cluster during growth in oxacillin (50). Furthermore oxacillin-induced repression of the *pur* operon was accompanied by significantly increased expression of *purR* (50), a finding in keeping with the subsequent reports that PurR also acts as a repressor of virulence (63, 64). Taken together, it appears that under β-lactam stress, MRSA may attempt to limit flux through purine biosynthetic pathways as a means of increasing resistance and that, conversely, increased guanosine or xanthosine flux into the GTP branch of purine metabolism has the reverse effect and reduces β-lactam resistance. A recent paper describing the interplay between *de novo* purine biosynthesis and c-di-AMP signalling demonstrated that a *purF* mutation was associated with reduced levels of c-di-AMP (35). Here we report that, with the exception of *purH*, mutations in all other *pur* operon genes including *purF* were accompanied by increased β-lactam resistance. The reduction in c-di-AMP levels in a *purF* mutant reported by Li et al (35) and the increase in oxacillin resistance described in this study may at first appear to be contradictory. However, it is important to note that the contribution of purine homeostasis to β-lactam resistance can only be accurately characterised when MRSA is grown in the presence of these antibiotics. Thus under standard growth conditions (MHB NaCl media), our data showed that the purine salvage pathway *nupG* and *deoD2* mutations were associated with reduced levels of c-di-AMP, whereas under very mild oxacillin stress, c-di-AMP was restored to wild-type levels in these mutants. Furthermore the reduced levels of c-di-AMP in wild-type exposed to the guanosine/oxacillin combination was reversed in the *nupG* and *deoD2* mutants.

Among the purine metabolism mutations examined, *purH* and *deoD1* were not found to have any impact on β-lactam resistance, although both mutants remained susceptible to OX/Gua treatment. Purine nucleoside phosphorylases typically form oligomers raising the possibility that DeoD1 and DeoD2 can heterodimerise, which may influence their contribution to purine salvage and β-lactam resistance, particularly if they are differentially regulated under specific growth conditions. The *deoD1* gene is co-located on the chromosome with a major facilitator superfamily (Mfs) gene, *deoC1* (deoxyribose-phosphate aldolase) and *deoB* (phosphopentomutase), which may be expressed as an operon. Mutations in the Mfs gene, *deoC1* and *deoB* genes alone were not associated with altered OX susceptibility (65). Nevertheless, DeoD1 activity may be dependent on the Mfs protein or a solute/ion transported by this membrane protein. In contrast the *deoD2* gene is monocistronic and a second deoxyribose-phosphate aldolase, *deoC2*, sharing 98% identity with *deoC1*, is transcribed divergently from *deoD2*. The *purH* gene, which encodes the bifunctional phosphoribosylaminoimidazolecarboxamide formyltransferase / IMP cyclohydrolase is the penultimate gene in the *purEKCSQLFMNHD* operon followed only by *purD*. A *purD* mutation, like all other *pur* operon mutations is accompanied by increased OX resistance. PurH is responsible for the final step in the purine biosynthetic pathway, converting 5-aminoimidazole-4-carboxamide ribonucleotide (AICAR) into IMP. AICAR is produced by PurB, which is also part of the purine salvage pathway and *purB* can be transcriptionally regulated independent of the *pur* operon (66). Unlike other Pur enzymes, the bifunctional activity of PurH (synthesis and break-down of IMP), combined with its requirement for AICAR from the *de novo* or purine salvage pathways points to a complex role of this enzyme in purine metabolism and may suggest that its impact on OX resistance is more dependent on growth conditions than other *pur* operon enzymes.

Arising from studies to uncover the therapeutic potential of bactericidal and bacteriostatic compounds produced by coagulase-negative staphylococci (CoNS) to compete with *S. aureus* in shared ecological niche, the Heinrich’s group identified the CoNS-produced purine analogue 6-thioguanine (6-TG) and revealed its potential to both inhibit growth of *S. aureus* by interfering with ribosome biogenesis and downregulate *agr*-controlled virulence under purine limited growth conditions (67). Resistance to 6-TG can arise through mutations in PbuG (also named StgP, 6-thioguanine permease) and the Hpt hypoxanthine phosphoribosyltransferase (53), demonstrating that guanine/6-TG are transported and metabolised by PbuG/StgP and Hpt, respectively. We showed here that guanosine was unable to resensitize the 6-TG resistant *hpt* mutants R10 and R11 (53) to OX, further supporting the role for Hpt in guanine/6-TG metabolism. Our screen of suppressor mutants resistant to OX/Gua combinations also identified stop codon mutations in *hpt* as well as *nupG, prs* and in transcriptional (*rpoB*) and translational (tRNAs, ClpX) machinery genes. Interestingly, while the R10 and R11 *hpt* mutants were resistant to guanosine/OX combinations, they were susceptible to OX/Xan presumably because xanthosine is processed by the Xpt phosphoribosyltransferase thus bypassing Hpt. On the other hand, the GS8 and GS25 OX/Gua suppressor mutants carrying a Hpt L_113_Stop mutation unexpectedly remained resistant to OX/Xan. Although the reason for this difference is unclear, one possibility is that Hpt also plays a role in processing xanthosine. If this is true, the Hpt amino acid substitutions in R10 and R11 must interfere with guanosine processing, but may not affect processing of xanthosine whereas the L_113_Stop mutation presumably blocks processing of both nucleosides. Interestingly, a M_1_I mutation in *hpt* likely to negatively impact translation initiation was previously implicated the HoR phenotype (44). Similarly a *guaA* deletion mutation resulted in HoR resistance. There are no transposon insertions in *hpt* or *guaA* in the NTML indicating that these genes are important for growth under standard laboratory conditions. Collectively our data are in keeping with previous studies indicating that mutations negatively impacting GTP biosynthetic pathway enzyme activity are accompanied by increased resistance to β-lactams and β-lactam/nucleoside combinations.

Our data showed that *nupG* and *deoD2* mutants significantly increased β-lactam resistance independent of changes in PBP2a levels. Several lines of evidence support the conclusion that purine homeostasis regulated β-lactam resistance is mechanistically separate from PBP2a expression. For instance, our data showed that guanosine also significantly reduces the OX MICs of a laboratory MSSA strain and the NTML *mecA* transposon mutant. Two *clpX* stop codon mutations (G_98_* and G_278_*) were also identified giving rise to resistance to OX/Gua and OX/Xan combinations, and very high OX MICs. The ClpXP chaperone-protease complex is known to control β-lactam resistance in MRSA (25, 26, 68), independent of changes in *mecA* transcription or PBP2a levels (25). Here we also identified RpoB T_480_K and T_622_I substitutions implicated in resistance to nucleoside/OX combinations. Two independent studies of *rpoB* and *rpoC* mutations associated with high-level β-lactam resistance ruled out changes in PBP2a expression (6, 62). In a recent analysis Giulieri et al (69) implicated both *gdpP*/*dacA*-regulated c-di-AMP signal transduction, as well as the ClpXP chaperone-protease complex in *mecA*-independent, low level OX resistance. Collectively these findings support the conclusion that altered regulation of PBP2a expression does not play an important role in the regulation of β-lactam susceptibility by nucleoside/OX combinations. Instead our data showing that guanosine alone, and in particular in combination with OX, significantly reduces c-di-AMP levels in a NupG/DeoD2-dependent manner and dramatically increases cell size. Osmoregulation and cell size are among the important phenotypes controlled by c-di-AMP (23, 33) suggesting that deregulation of cellular osmotic control by exogenous purine nucleosides is the most likely mechanism for increased β-lactam susceptibility. From a therapeutic perspective, our *in vitro* kill curves revealed significant bactericidal activity using combinations guanosine and OX at concentrations up to 128 μg/ml, which are within the range of flucloxacillin intravenously administered to human patients (100-200 mg/kg/day). While the emergence of OX/Gua suppressor mutants appeared to be significantly reduced at higher oxacillin concentrations (as evidenced by the absence of re-growth in kill curve cultures), this remains a potential barrier to the use of nucleosides as β-lactam adjuvants. Future studies to measure the anti-MRSA activity of different purine nucleosides or purine analogues in combination with different β-lactams may identify more effective therapeutic regimes. Nevertheless, drugs derived from purine and pyrimidine nucleotides are already used for the treatment of cancer and viral infections highlighting the potential clinical applications of these findings, not only for MRSA but also other pathogens on the World Health Organisation’s list of priority antibiotic-resistant pathogens. Using purine nucleosides as adjuvants to increase the susceptibility of clinically important pathogens to β-lactams has the potential to facilitate new treatments with lower antibiotic doses and with drug combinations that are toxic at higher concentrations.

## Materials and Methods

### Bacterial strains and growth conditions

Bacterial strains and plasmids used in this study are listed in Table S1. *Escherichia coli* strains were grown in Luria Bertani (LB) broth or agar (LBA) and *Staphylococcus aureus* strains were grown in Mueller-Hinton Broth (MHB), Mueller-Hinton Agar (MHA), Tryptic Soy Broth (TSB), Tryptic Soy Agar (TSA), chemically defined medium (CDM), supplemented with erythromycin (Erm) 10 μg/ml, chloramphenicol (Cm) 10 μg/ml, ampicillin (Amp) 100 μg/ml, kanamycin (Km) 90 μg/ml, guanosine, xanthosine or adenosine (0.2 – 1 g/l) where indicated.

Data from growth experiments in a Tecan Sunrise microplate instrument were recorded using Magellan software. Overnight cultures were grown in 5 ml TSB using a single colony, and were washed once in PBS before being used to inoculate CDM cultures. PBS washed overnight cultures were adjusted to an OD_600_ of 0.1 in PBS and 10 μl inoculated into 200 μl media per well before being incubated at 35°C for 24 h with shaking and *A*_600_ recorded every 15 min intervals.

For flask growth curves, 250 ml flasks were filled with 25 ml growth media, and cultures grown overnight in TSB were used to inoculate the media at a starting *A*_600_ of 0.01. Flasks were incubated at 37°C shaking at 200 rpm. *A*_600_ readings were measured at 2 h intervals for 12 hours and an extra reading was taken at 24 hours.

Colony forming units (CFUs) were enumerated by serially diluting 100 μl aliquots removed from flask cultures. Plates were incubated for 20 h at 37 °C and colonies enumerated, and CFUs per ml suspension per OD_600_ calculated. Three independent biological replicates were performed for each strain and the resulting data plotted using GraphPad Prism software.

### Genetic manipulation of *S. aureus*

Phage 80α was used as described previously (70) to transduce the transposon insertions from NE650, NE1419 into the parent JE2 strain to rule out the possibility that uncharacterised mutations in the NTML mutants may contribute to the increased antibiotic resistance phenotypes. Transposon insertions were verified by PCR amplification of the target loci using primers listed in Table S2.

To complement NE650 and NE1419, 1,387bp and 1,740bp PCR fragments were amplified using JE2 gDNA as template and the primers INF_deoD_F / INF_deoD_R and INF_nupG_F / INF_nupG_R, respectively (Table S2, Fig S4). The Clontech Infusion Cloning kit was used to insert the fragments into the *Eco*R1 restriction site of pLI50 to generate pLI50_*nupG*, and pLI50_*deoD2*. The recombinant plasmids were first transformed into cold-competent *E. coli* HST08 before being transformed by electroporation into the restriction-deficient *S. aureus* strain RN4220 and finally into NE650 and NE1419.

### PBP2a Western blot analysis

Single colonies from wild-type JE2, NE1419 (*nupG*), NE650 (*deoD2*), NE1419 pLI50_*nupG*, NE650 pLI50_*deoD2*, MSSA strain 8325-4 (negative control) and HoR MRSA strain BH1CC (positive control) were inoculated in MHB 2% NaCl overnight and grown at 35°C with 200 rpm shaking. All cultures were supplemented with 5 μg/ml OX, with the exception of 8325-4 which was grown in MHB 2% NaCl without antibiotic and BHICC which was grown with 32 μg/ml OX. The next day, samples were pelleted and resuspended in PBS to an *A*_600_ = 10. Six μl of lysostaphin (10 μg/ml) and 1 μl of DNase (10 µg/ml) was added to 500 μl of this concentrated cell suspension before being incubated at 37°C for 40 min. Next, 50 μl of 10% SDS was added and the incubation continued for a further 20 min. The lysed cells were then pelleted in a microcentrifuge for 15 min, following which the protein-containing supernatant was collected and total protein concentration determined using the Pierce BCA Protein Assay Kit. For each sample, 6 μg total protein was run on a 7.5% Tris-Glycine gel, transferred to a PVDF membrane and probed with anti-PBP2a (1:1000), followed by HRP-conjugated protein G (1:2000) and colorimetric detection with Opti-4CN Substrate kit. Three independent experiments were performed.

### Antibiotic disc diffusion susceptibility assays

Disk diffusion susceptibility testing was performed in accordance with Clinical Laboratory Standards Institute (CLSI) guidelines (71) with the following modifications. Overnight cultures were diluted into fresh TSB and grown for 3 h at 37 °C. Day cultures were then adjusted to *A*_600_ = 0.5 and 150 μl of this suspension was swabbed evenly 3 times across the surface of an MHA plate (4 mm agar depth). 6 mm antibiotic discs were applied, before incubation for times specified by CLSI guidelines for stated antibiotics at 35 - 37°C. Twenty μl of guanosine at varying concentrations (50 - 750 μg/ml) was spotted onto discs to demonstrate synergistic activity with β-lactam antibiotics where indicated. Three independent measurements were performed for each strain.

### Antibiotic minimum inhibitory concentration (MIC) and synergy assays

MIC measurements by broth microdilutions were performed in accordance with CLSI methods for dilution susceptibility testing of staphylococci (72) with modifications as follows: guanosine, xanthosine or adenosine were supplemented into culture media at concentrations ranging from 200 – 1000 μg/ml. Strains were first grown at 37 °C on MHA (or MHB NaCl for OX MIC assays) for 24 h and 5 - 6 colonies were resuspended in 0.85% saline before being adjusted to 0.5 McFarland standard (*A*_600_ = 0.01). The cell suspension was then diluted 1:20 in PBS and 20 μl used to inoculate 180 μl MHB / MHB NaCl supplemented with purines as indicated in the individual wells of 96-well plates. The plates were incubated at 37 °C for 24 h and MIC values were recorded as the lowest antibiotic concentration where no growth was observed.

### Isolation of suppressor mutants resistant to guanosine/oxacillin combination

Suppressor mutants were isolated from JE2 cultures serially passaged in MHB 2% NaCl supplemented with 200 μg/ml guanosine and 32 μg/ml OX at 35°C. Using 96-well plates, 100 μl of culture medium was inoculated with 2 - 8 × 10^5^ CFU/ml from a JE2 overnight culture and growth was monitored for 24 h in a Tecan Sunrise microplate instrument using Magellan software, after which a 1:20 dilution was used to inoculate 100 μl of the same medium in the corresponding well of a new plate, which was incubated for a further 24 h. In wells showing visible growth, 5 μl of culture was inoculated onto TSA plates to purify single colonies from the suppressor strains. The susceptibility (MICs) of each of the isolated suppressor strains to OX and OX/GUA combinations was confirmed as described above.

### Genomic DNA (gDNA) extraction and whole-genome sequencing (WGS)

Genomic DNA (gDNA) extractions were performed using the Wizard® Genomic DNA Purification Kit (Promega) following pre-treatment of *S. aureus* cells with 10 μg/ml lysostaphin (Ambi Products LLC) at 37°C for 30 min. DNA libraries were prepared using an Illumina Nextera DNA kit. An Illumina MiSeq v2 300 cycle kit was used for the genome sequencing and the 150 bp paired end sequencing run was performed at the MRC London Institute of Medical Sciences Genomics Facility. The CLC Genomics Workbench software (Qiagen Version 20) was used for genome sequencing analysis of the different strains, as described previously (39). As a reference genome, a contig was produced for wild-type JE2 by mapping Illumina reads onto the closely related USA300 FPR3757 genome sequence (RefSeq accession number NC_07793.1). The Illumina short read sequences from NTML mutants of interest were then mapped onto the assembled JE2 sequence, and the presence of the transposon insertion confirmed. Single Nucleotide Polymorphisms (SNPs), deletions or insertions were mapped to the genomes of the NTML mutants and OX/Gua resistant suppressor strains and the presence of large deletions ruled out by manually searching for zero coverage regions. Whole genome sequence data is available from the European Nucleotide Archive (Study Accession number PRJEB55671), accession numbers ERS13358424 - ERS13358446).

### Antibiotic kill curves

Kill Curves were performed according to CLSI guidelines to determine the extent of synergy between purines and oxacillin. Overnight cultures grown in MHB NaCl were diluted 1:100 and allowed to grow for three hours (exponential growth phase) before being inoculated into 25 ml of MHB NaCl in 250 ml flasks at a starting cell density of approximately 1×10^6^ CFU/ml. The flasks were incubated at 35°C with shaking at 200 rpm and CFUs were enumerated after 0, 2, 4, 6, 8, 12 and 24 h by serially diluting and plating 100 μl aliquots removed from flask cultures on tryptone soya agar plates.

### Microscopy and cell size determination

For microscopy experiments, MRSA strain JE2 was grown overnight at 37°C in MHB. The next day, it was back-diluted into MHB, MHB Gua 0.2 g/l, MHB OX 1 μg/ml and MHB Gua 0.2 g/l OX 1 μg/ml and day cultures were grown for 5 hours at 35°C. After the 5-hour growth, the day cultures were washed and normalized to an *A*_600_ of 1 in PBS and 75 μl of these cultures were double stained for 30 mins at 37 °C with vancomycin-BODIPY FL at a final concentration of 2 μg/ml and WGA Alexa Fluor 594 at a final concentration of 25 μg/ml. Bacteria were then collected by centrifugation for 2 mins at 14,000 x*g*. The cells were resuspended with 100 μl of PBS, pH 7.4, and 5 μl of this sample was spotted onto a thin 1% agarose gel patch prepared in PBS. Stained bacteria were then imaged at X1000 magnification using Olympus LS FLUOVIEW Fv3000 Confocal Laser Scanning Microscope. Cell size was measured as previously described (39) using ImageJ software. Images of cells from three biological replicates were acquired, 40 cells measured per biological replicate (120 cells in total per condition), and the average and standard deviations for the three biological replicates were plotted using GraphPad Prism version 9.2 and significant differences were determined using one-way ANOVA followed by Tukey’s post-hoc.

### GTP assays

A GTPase-Glo bioluminescence assay kit (Promega) was used and the manufacturer’s guidelines were adjusted to measure relative intracellular GTP levels by luminescence. Briefly overnight cultures of JE2, *nupG* and *deoD2* grown at 37°C in MHB NaCl were used to inoculate duplicate MHB NaCl cultures supplemented with guanosine (Gua, 0.2 g/l) and/or OX (OX, 1 μg/ml) at a starting *A*_600_ = 0.05 before these were again incubated overnight at 37°C. Cell pellets were lysed and total protein concentrations determined using a Bicinchoninic Acid (BCA) assay. The protein concentration in each lysate was adjusted to 50 μg/ml and 2.5 μl (125 ng) used in each GTPase-Glo assay.

### Measurement of (p)ppGpp

Cultures of JE2, *nupG* and *deoD2* were grown overnight at 37°C in MHB, followed by dilution in MHB NaCl with OX (1 μg/ml) and/or guanosine (0.2 g/l) to *A*_600_ = 0.05. Cells were grown to *A*_600_ = 0.5 and divided into 2 aliquots. To one, 3.7 MBq of (^32^P)H_3_PO_4_ was added, while the other lacked radiation to allow for normalization of cultures via protein concentrations by BCA assay. Both sets of culture were grown overnight and the non-radioactive set used to determine protein concentration. Radioactive culture corresponding to 150 ng of protein were lysed by the addition of 2 M formic acid and subsequently subjected to four freeze/thaw cycles. Ten μl of the supernatant fractions were subsequently spotted on PEI-cellulose F TLC plates (Merck Millipore) and nucleotides separated using a 1.5 M KH_2_PO_4_, pH 3.6, buffer. The radioactive spots were visualized using an FLA 7000 Typhoon PhosphorImager, and data were quantified using ImageQuantTL software.

### Quantification of c-di-AMP levels using a competitive ELISA

Relative intracellular levels of c-di-AMP were determined using a previously described competitive ELISA method (73) with some modifications. Briefly, single colonies of wild-type JE2, *nupG* (NE1419) and *deoD2* (NE650) were picked from TSA plates and used to inoculate 5 ml or 12 ml of MHB 2% NaCl with or without guanosine (0.2 g/l) and/or OX (0.2 μg/ml) and the cultures were incubated for 18 h at 37°C with shaking. The 5ml cultures were grown in 20 ml tubes and 12 ml cultures were grown in 125 ml flasks. Next day, 1.5 ml (for all strains grown without OX), 3 ml (for *nupG* and *deoD2* grown in OX and the wild-type grown in OX alone) or 6 ml (for wild-type JE2 grown in both OX and guanosine) were collected by centrifugation, washed three times with PBS, resuspended in 0.75 ml of 50 mM Tris pH 8 buffer supplemented with 20 ng/ml lysostaphin and incubated for 1 h before completing cell lysis by bead beating. Following centrifugation for 5 min at 17,000 ×*g*, the supernatant from the cell lysates were transferred to new tubes and protein concentration of the samples was determined using a Pierce BCA protein assay kit (Thermo Scientific, Waltham, MA, USA). The samples were then heated to 95°C for 10 min, centrifuged for 5 min at 17,000 ×*g* and the supernatant containing c-di-AMP was transferred to a new tube. The samples were diluted to a protein concentration of 300-800 μg/ml, as appropriate. 100 μl of coating buffer (50 mM Na_2_C0_3_, 50 mM NaHCO_3_, pH 9.6) containing 10 μg/ml of the c-di-AMP binding protein CpaA_SP_ was aliquoted into each well of a 96 well NUNC MaxiSorp plate (Thermo Scientific, Waltham, MA, USA), which was then incubated for 18 h at 4°C. The plate was then washed three times with 200 μl PBST pH 7.4 (10 mM Na_2_HPO_4_, 1.8 mM KH_2_PO_4_ 137 mM NaCl, 2.7 mM KCl, 0.05% (v/v) Tween 20), blocked for 1 h at 18°C with 150 μl blocking solution (1% BSA in PBS pH 7.4) and again washed three times with 200 μl PBST. 250 μl aliquots of the samples (with three technical replicates for each biological replicate) or standards (two technical replicates) were then mixed with 250 μl of a 50 nM biotinylated c-di-AMP solution prepared in 50 mM Tris pH 8 buffer. For the standard curve, c-di-AMP standards were prepared in 50 mM Tris pH 8 buffer at concentrations of 0, 12.5, 25, 37.5, 50, 75, 100 and 200 nM. Following the addition of the samples and the standards, the 96 well plates were incubated for 2 h at 18°C and then washed three times with PBST. Next, 100 μl of a high-sensitivity streptavidin-HRP solution (Thermo Scientific, Waltham, MA, USA) diluted 1:500 in PBS was added to each well and the plate was incubated for 1 h at 18°C. The plate was washed again three times with 200 μl PBST and 100 μl of a developing solution (0.103 M NaHPO_4_, 0.0485 M citric acid, 500 mg/L o-phenylenediamine dihydrochloride, 0.03% H_2_O_2_) was added to each well and the plate incubated for 15 min at 18°C. The reaction was stopped by adding 100 μl of 2 M H_2_SO_4_ solution and the absorbance measured in a plate reader at 490 nm. c-di-AMP concentrations were calculated as ng c-di-AMP/mg protein. The data from three independent experiments was collated and presented by showing relative c-di-AMP levels in all samples compared to the wild-type grown in MHB NaCl, MHG NaCl Ox or MHB NaCl Ox/Gua as appropriate, which was assigned a value of 100%. Standard deviations are shown and significant differences were determined using one-way ANOVA followed by Tukey’s post-hoc tests.

## Data availability

Whole genome sequence data is available from the European Nucleotide Archive (Study Accession number PRJEB55671), accession numbers ERS13358424 - ERS13358446. The SAUSA300_FRP3757 (TaxID:451515) reference genome sequence is available from NCBI.

## Author contributions

Aaron Nolan: Conceptualization, Formal analysis, Investigation, Methodology, Writing - original draft, Writing - review & editing

Merve S. Zeden: Conceptualization, Formal analysis, Investigation, Data curation, Methodology, Supervision, Writing - original draft, Writing - review & editing

Christopher Campbell: Conceptualization, Formal analysis, Investigation, Methodology, Writing - review & editing

Igor Kviatkovski: Formal analysis, Investigation, Methodology, Writing - review & editing

Lucy Urwin: Formal analysis, Investigation, Methodology, Writing - review & editing

Rebecca M. Corrigan: Funding acquisition, Formal analysis, Investigation, Methodology, Supervision, Writing - review & editing

Angelika Gründling: Funding acquisition, Formal analysis, Investigation, Methodology, Supervision, Writing - review & editing

James P. O’Gara: Conceptualization, Formal analysis, Funding acquisition, Project administration, Supervision, Writing - original draft, Writing - review & editing

## Acknowledgements

A.N., M.S.Z., C.C. and J.P.O’G. were supported by grants from the Health Research Board (ILP-POR-2019-102) (www.hrb.ie), the Irish Research Council (GOIPG/2014/763) (www.research.ie) and Science Foundation Ireland (19/FFP/6441) (www.sfi.ie). L.U. and R.M.C. were supported by a Sir Henry Dale Fellowship jointly funded by the Wellcome Trust and the Royal Society (104110/Z/14/A). I.K. and A.G. were supported by Wellcome trust grant 210671/Z/18/Z/WT. We are grateful to Prof. David Heinrichs for the gift of strains *S. aureus* R10 and R11 and Dr. Peter Owens from the Centre for Microscopy & Imaging at the University of Galway (www.imaging.nuigalway.ie) for his scientific and technical assistance. The funders had no role in study design, data collection and interpretation, or the decision to submit the work for publication.

## Supplementary Tables

**Table S1.**
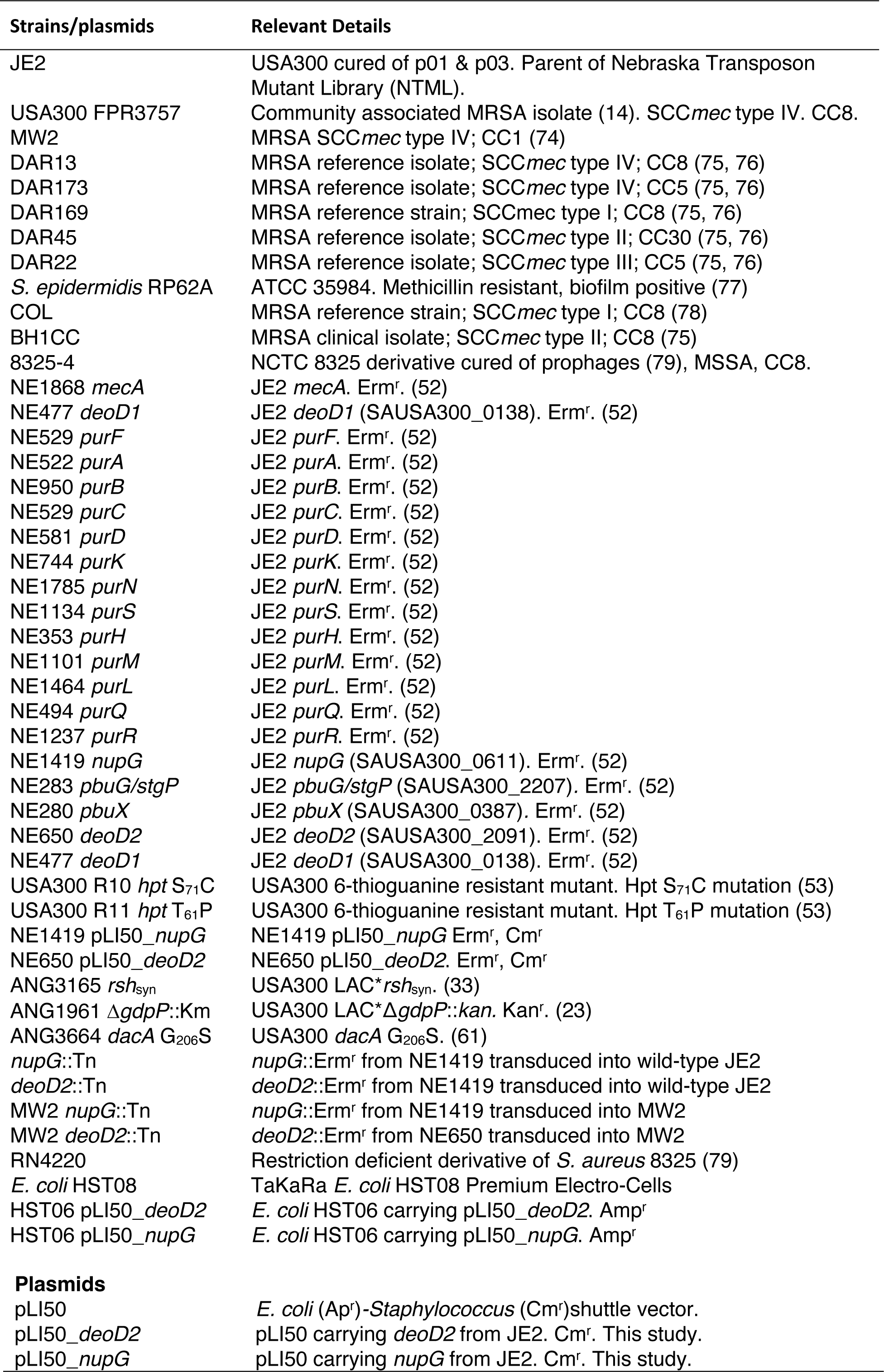
Bacterial strains and plasmids used in this study

| Strains/plasmids | Relevant Details |
| --- | --- |
| JE2 | USA300 cured of p01 & p03. Parent of Nebraska Transposon Mutant Library (NTML). |
| USA300 FPR3757 | Community associated MRSA isolate (14). SCCmec type IV. CC8. |
| MW2 | MRSA SCCmec type IV; CC1 (74) |
| DAR13 | MRSA reference isolate; SCCmec type IV; CC8 (75, 76) |
| DAR173 | MRSA reference isolate; SCCmec type IV; CC5 (75, 76) |
| DAR169 | MRSA reference strain; SCCmec type I; CC8 (75, 76) |
| DAR45 | MRSA reference isolate; SCCmec type II; CC30 (75, 76) |
| DAR22 | MRSA reference isolate; SCCmec type III; CC5 (75, 76) |
| *S. epidermidis* RP62A | ATCC 35984. Methicillin resistant, biofilm positive (77) |
| COL | MRSA reference strain; SCCmec type I; CC8 (78) |
| BH1CC | MRSA clinical isolate; SCCmec type II; CC8 (75) |
| 8325-4 | NCTC 8325 derivative cured of prophages (79), MSSA, CC8. |
| NE1868 *mecA* | JE2 *mecA*. Erm<sup>r</sup>. (52) |
| NE477 *deoD1* | JE2 *deoD1* (SAUSA300_0138). Erm<sup>r</sup>. (52) |
| NE529 *purF* | JE2 *purF*. Erm<sup>r</sup>. (52) |
| NE522 *purA* | JE2 *purA*. Erm<sup>r</sup>. (52) |
| NE950 *purB* | JE2 *purB*. Erm<sup>r</sup>. (52) |
| NE529 *purC* | JE2 *purC*. Erm<sup>r</sup>. (52) |
| NE581 *purD* | JE2 *purD*. Erm<sup>r</sup>. (52) |
| NE744 *purK* | JE2 *purK*. Erm<sup>r</sup>. (52) |
| NE1785 *purN* | JE2 *purN*. Erm<sup>r</sup>. (52) |
| NE1134 *purS* | JE2 *purS*. Erm<sup>r</sup>. (52) |
| NE353 *purH* | JE2 *purH*. Erm<sup>r</sup>. (52) |
| NE1101 *purM* | JE2 *purM*. Erm<sup>r</sup>. (52) |
| NE1464 *purL* | JE2 *purL*. Erm<sup>r</sup>. (52) |
| NE494 *purQ* | JE2 *purQ*. Erm<sup>r</sup>. (52) |
| NE1237 *purR* | JE2 *purR*. Erm<sup>r</sup>. (52) |
| NE1419 *nupG* | JE2 *nupG* (SAUSA300_0611). Erm<sup>r</sup>. (52) |
| NE283 *pbuG/stgP* | JE2 *pbuG/stgP* (SAUSA300_2207). Erm<sup>r</sup>. (52) |
| NE280 *pbuX* | JE2 *pbuX* (SAUSA300_0387). Erm<sup>r</sup>. (52) |
| NE650 *deoD2* | JE2 *deoD2* (SAUSA300_2091). Erm<sup>r</sup>. (52) |
| NE477 *deoD1* | JE2 *deoD1* (SAUSA300_0138). Erm<sup>r</sup>. (52) |
| USA300 R10 *hpt* S<sub>71</sub>C | USA300 6-thioguanine resistant mutant. Hpt S<sub>71</sub>C mutation (53) |
| USA300 R11 *hpt* T<sub>61</sub>P | USA300 6-thioguanine resistant mutant. Hpt T<sub>61</sub>P mutation (53) |
| NE1419 pLI50_ *nupG* | NE1419 pLI50_ *nupG* Erm<sup>r</sup>, Cm<sup>r</sup> |
| NE650 pLI50_ *deoD2* | NE650 pLI50_ *deoD2*. Erm<sup>r</sup>, Cm<sup>r</sup> |
| ANG3165 *rsh*<sub>syn</sub> | USA300 LAC* *rsh* <sub>syn</sub>. (33) |
| ANG1961 Δ*gdpP*::Km | USA300 LAC*Δ*gdpP*::*kan*. Kan<sup>r</sup>. (23) |
| ANG3664 *dacA* G<sub>206</sub>S | USA300 *dacA* G<sub>206</sub>S. (61) |
| *nupG*::Tn | *nupG*::Erm<sup>r</sup> from NE1419 transduced into wild-type JE2 |
| *deoD2*::Tn | *deoD2*::Erm<sup>r</sup> from NE1419 transduced into wild-type JE2 |
| MW2 *nupG*::Tn | *nupG*::Erm<sup>r</sup> from NE1419 transduced into MW2 |
| MW2 *deoD2*::Tn | *deoD2*::Erm<sup>r</sup> from NE650 transduced into MW2 |
| RN4220 | Restriction deficient derivative of *S. aureus* 8325 (79) |
| *E. coli* HST08 | TaKaRa *E. coli* HST08 Premium Electro-Cells |
| HST06 pLI50_ *deoD2* | *E. coli* HST06 carrying pLI50_ *deoD2*. Amp<sup>r</sup> |
| HST06 pLI50_ *nupG* | *E. coli* HST06 carrying pLI50_ *nupG*. Amp<sup>r</sup> |
| Plasmids |  |
| pLI50 | *E. coli* (Ap<sup>r</sup>)-*Staphylococcus* (Cm<sup>r</sup>)shuttle vector. |
| pLI50_ *deoD2* | pLI50 carrying *deoD2* from JE2. Cm<sup>r</sup>. This study. |
| pLI50_ *nupG* | pLI50 carrying *nupG* from JE2. Cm<sup>r</sup>. This study. |

**Table S2.**
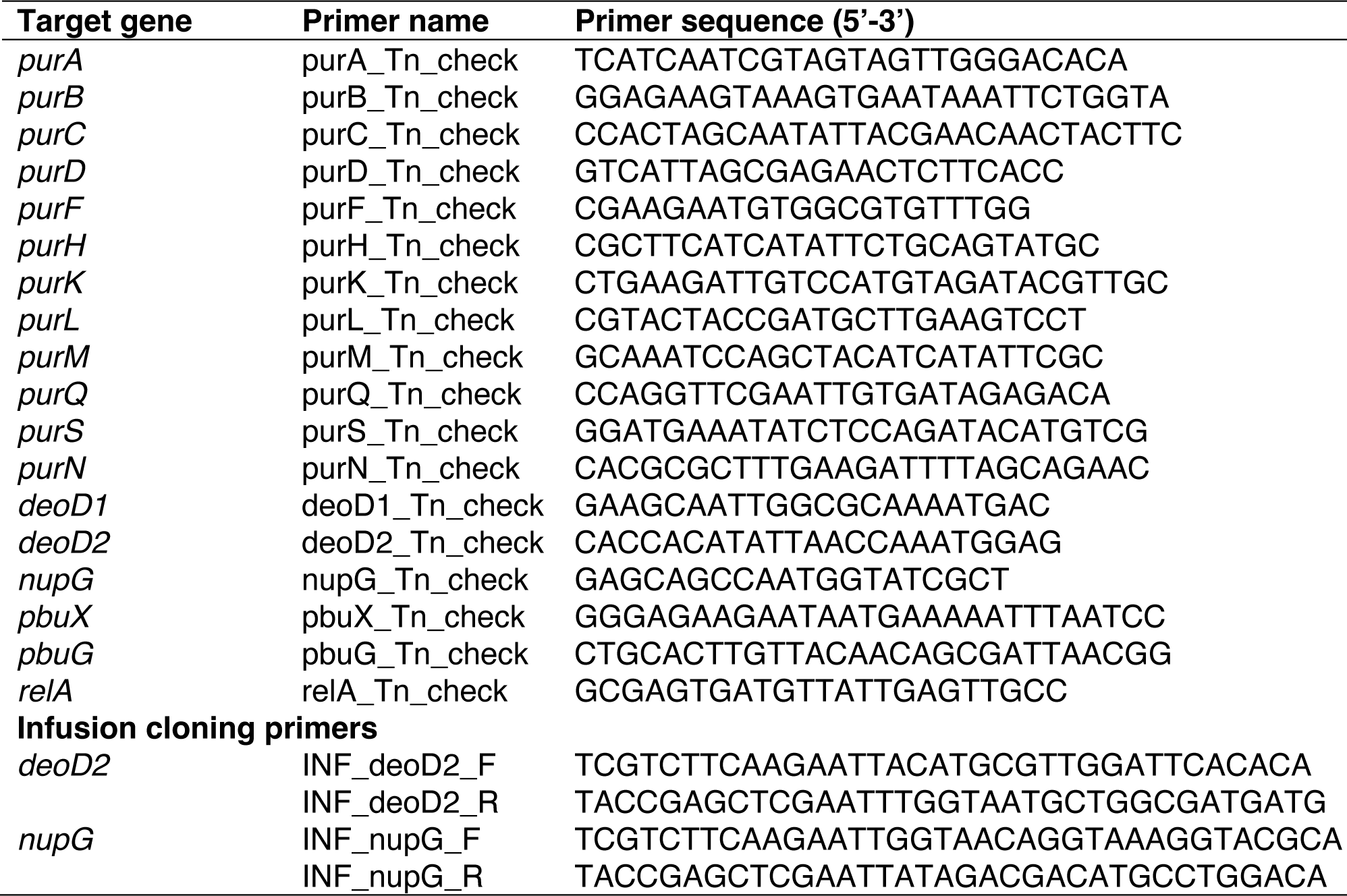
Oligonucleotide primers used in this study.

## Notes

### Competing Interest Statement

The authors have declared no competing interest.

### Summary of Updates

Supplemental files added.

## References

1. Lowy FD. 1998. Staphylococcus aureus infections. N Engl J Med 339:520–32.

2. Martins A, Cunha Mde L. 2007. Methicillin resistance in Staphylococcus aureus and coagulase-negative staphylococci: epidemiological and molecular aspects. Microbiol Immunol 51:787–95.

3. van Hal SJ, Jensen SO, Vaska VL, Espedido BA, Paterson DL, Gosbell IB. 2012. Predictors of mortality in Staphylococcus aureus Bacteremia. Clin Microbiol Rev 25:362–86.

4. Liu C, Bayer A, Cosgrove SE, Daum RS, Fridkin SK, Gorwitz RJ, Kaplan SL, Karchmer AW, Levine DP, Murray BE, M JR, Talan DA, Chambers HF. 2011. Clinical practice guidelines by the infectious diseases society of america for the treatment of methicillin-resistant Staphylococcus aureus infections in adults and children: executive summary. Clin Infect Dis 52:285–92.

5. Hiramatsu K, Ito T, Tsubakishita S, Sasaki T, Takeuchi F, Morimoto Y, Katayama Y, Matsuo M, Kuwahara-Arai K, Hishinuma T, Baba T. 2013. Genomic Basis for Methicillin Resistance in Staphylococcus aureus. Infect Chemother 45:117–36.

6. Panchal VV, Griffiths C, Mosaei H, Bilyk B, Sutton JAF, Carnell OT, Hornby DP, Green J, Hobbs JK, Kelley WL, Zenkin N, Foster SJ. 2020. Evolving MRSA: High-level beta-lactam resistance in Staphylococcus aureus is associated with RNA Polymerase alterations and fine tuning of gene expression. PLoS Pathog 16:e1008672.

7. Bruniera FR, Ferreira FM, Saviolli LR, Bacci MR, Feder D, da Luz Gonçalves Pedreira M, Sorgini Peterlini MA, Azzalis LA, Campos Junqueira VB, Fonseca FL. 2015. The use of vancomycin with its therapeutic and adverse effects: a review. Eur Rev Med Pharmacol Sci 19:694–700.

8. Rybak MJ, Lomaestro BM, Rotscahfer JC, Moellering RC, Jr., Craig WA, Billeter M, Dalovisio JR, Levine DP. 2009. Vancomycin Therapeutic Guidelines: A Summary of Consensus Recommendations from the Infectious Diseases Society of America, the American Society of Health-System Pharmacists, and the Society of Infectious Diseases Pharmacists. Clinical Infectious Diseases 49:325–327.

9. Lamp KC, Rybak MJ, Bailey EM, Kaatz GW. 1992. In vitro pharmacodynamic effects of concentration, pH, and growth phase on serum bactericidal activities of daptomycin and vancomycin. Antimicrobial agents and chemotherapy 36:2709–2714.

10. Howe RA, Monk A, Wootton M, Walsh TR, Enright MC. 2004. Vancomycin susceptibility within methicillin-resistant Staphylococcus aureus lineages. Emerging infectious diseases 10:855–857.

11. Mwangi MM, Wu SW, Zhou Y, Sieradzki K, de Lencastre H, Richardson P, Bruce D, Rubin E, Myers E, Siggia ED, Tomasz A. 2007. Tracking the in vivo evolution of multidrug resistance in Staphylococcus aureus by whole-genome sequencing. Proceedings of the National Academy of Sciences of the United States of America 104:9451–9456.

12. Chambers HF. 2001. Methicillin-resistant Staphylococcus aureus. Mechanisms of resistance and implications for treatment. Postgrad Med 109:43–50.

13. Hartman BJ, Tomasz A. 1984. Low-affinity penicillin-binding protein associated with beta-lactam resistance in Staphylococcus aureus. J Bacteriol 158:513–6.

14. Diep BA, Gill SR, Chang RF, Phan TH, Chen JH, Davidson MG, Lin F, Lin J, Carleton HA, Mongodin EF, Sensabaugh GF, Perdreau-Remington F. 2006. Complete genome sequence of USA300, an epidemic clone of community-acquired meticillin-resistant Staphylococcus aureus. Lancet 367:731–9.

15. Francis JS, Doherty MC, Lopatin U, Johnston CP, Sinha G, Ross T, Cai M, Hansel NN, Perl T, Ticehurst JR, Carroll K, Thomas DL, Nuermberger E, Bartlett JG. 2005. Severe community-onset pneumonia in healthy adults caused by methicillin-resistant Staphylococcus aureus carrying the Panton-Valentine leukocidin genes. Clin Infect Dis 40:100–7.

16. Katayama Y, Ito T, Hiramatsu K. 2000. A new class of genetic element, staphylococcus cassette chromosome mec, encodes methicillin resistance in Staphylococcus aureus. Antimicrob Agents Chemother 44:1549–55.

17. Llarrull LI, Fisher JF, Mobashery S. 2009. Molecular basis and phenotype of methicillin resistance in Staphylococcus aureus and insights into new beta-lactams that meet the challenge. Antimicrob Agents Chemother 53:4051–63.

18. Finan JE, Rosato AE, Dickinson TM, Ko D, Archer GL. 2002. Conversion of oxacillin-resistant staphylococci from heterotypic to homotypic resistance expression. Antimicrob Agents Chemother 46:24–30.

19. Tomasz A, Nachman S, Leaf H. 1991. Stable classes of phenotypic expression in methicillin-resistant clinical isolates of staphylococci. Antimicrobial agents and chemotherapy 35:124–129.

20. Kim C, Mwangi M, Chung M, Milheirico C, de Lencastre H, Tomasz A. 2013. The mechanism of heterogeneous beta-lactam resistance in MRSA: key role of the stringent stress response. PLoS One 8:e82814.

21. Mwangi MM, Kim C, Chung M, Tsai J, Vijayadamodar G, Benitez M, Jarvie TP, D. L, Tomasz A. 2013. Whole-genome sequencing reveals a link between beta-lactam resistance and synthetases of the alarmone (p)ppGpp in Staphylococcus aureus. Microb Drug Resist 19:153–9.

22. Pozzi C, Waters EM, Rudkin JK, Schaeffer CR, Lohan AJ, Tong P, Loftus BJ, Pier GB, Fey PD, Massey RC, O’Gara JP. 2012. Methicillin resistance alters the biofilm phenotype and attenuates virulence in Staphylococcus aureus device-associated infections. PLoS Pathog 8:e1002626.

23. Corrigan RM, Abbott JC, Burhenne H, Kaever V, Gründling A. 2011. c-di-AMP is a new second messenger in Staphylococcus aureus with a role in controlling cell size and envelope stress. PLoS Pathog 7:e1002217.

24. Griffiths JM, O’Neill AJ. 2012. Loss of function of the gdpP protein leads to joint beta-lactam/glycopeptide tolerance in Staphylococcus aureus. Antimicrob Agents Chemother 56:579–81.

25. Baek KT, Gründling A, Mogensen RG, Thogersen L, Petersen A, Paulander W, Frees D. 2014. beta-Lactam resistance in methicillin-resistant Staphylococcus aureus USA300 is increased by inactivation of the ClpXP protease. Antimicrob Agents Chemother 58:4593–603.

26. Jensen C, Baek KT, Gallay C, Thalso-Madsen I, Xu L, Jousselin A, Ruiz Torrubia F, Paulander W, Pereira AR, Veening JW, Pinho MG, Frees D. 2019. The ClpX chaperone controls autolytic splitting of Staphylococcus aureus daughter cells, but is bypassed by beta-lactam antibiotics or inhibitors of WTA biosynthesis. PLoS Pathog 15:e1008044.

27. Giuliani AL, Sarti AC, Di Virgilio F. 2019. Extracellular nucleotides and nucleosides as signalling molecules. Immunology Letters 205:16–24.

28. Kilstrup M, Hammer K, Ruhdal Jensen P, Martinussen J. 2005. Nucleotide metabolism and its control in lactic acid bacteria. FEMS Microbiology Reviews 29:555–590.

29. Pesavento C, Hengge R. 2009. Bacterial nucleotide-based second messengers. Current Opinion in Microbiology 12:170–176.

30. Gründling A, Lee VT. 2016. Old concepts, new molecules and current approaches applied to the bacterial nucleotide signalling field. Philosophical Transactions of the Royal Society B: Biological Sciences 371:20150503.

31. Dengler V, McCallum N, Kiefer P, Christen P, Patrignani A, Vorholt JA, Berger-Bachi B, Senn MM. 2013. Mutation in the C-di-AMP cyclase dacA affects fitness and resistance of methicillin resistant Staphylococcus aureus. PLoS One 8:e73512.

32. Fahmi T, Port GC, Cho KH. 2017. c-di-AMP: An Essential Molecule in the Signaling Pathways that Regulate the Viability and Virulence of Gram-Positive Bacteria. Genes (Basel) 8.

33. Corrigan RM, Bowman L, Willis AR, Kaever V, Gründling A. 2015. Cross-talk between two nucleotide-signaling pathways in Staphylococcus aureus. J Biol Chem 290:5826–39.

34. Li L, Abdelhady W, Donegan NP, Seidl K, Cheung A, Zhou Y-F, Yeaman MR, Bayer AS, Xiong YQ. 2018. Role of Purine Biosynthesis in Persistent Methicillin-Resistant Staphylococcus aureus Infection. The Journal of infectious diseases 218:1367–1377.

35. Li L, Li Y, Zhu F, Cheung AL, Wang G, Bai G, Proctor RA, Yeaman MR, Bayer AS, Xiong YQ. 2021. New Mechanistic Insights into Purine Biosynthesis with Second Messenger c-di-AMP in Relation to Biofilm-Related Persistent Methicillin-Resistant Staphylococcus aureus Infections. mBio 12:e0208121–e0208121.

36. Lan L, Cheng A, Dunman PM, Missiakas D, He C. 2010. Golden pigment production and virulence gene expression are affected by metabolisms in Staphylococcus aureus. J Bacteriol 192:3068–77.

37. Stulke J, Kruger L. 2020. Cyclic di-AMP Signaling in Bacteria. Annu Rev Microbiol 74:159–179.

38. Schuster CF, Bellows LE, Tosi T, Campeotto I, Corrigan RM, Freemont P, Gründling A. 2016. The second messenger c-di-AMP inhibits the osmolyte uptake system OpuC in Staphylococcus aureus. Sci Signal 9:ra81.

39. Zeden MS, Schuster CF, Bowman L, Zhong Q, Williams HD, Gründling A. 2018. Cyclic di-adenosine monophosphate (c-di-AMP) is required for osmotic regulation in Staphylococcus aureus but dispensable for viability in anaerobic conditions. J Biol Chem 293:3180–3200.

40. Corrigan RM, Gründling A. 2013. Cyclic di-AMP: another second messenger enters the fray. Nat Rev Microbiol 11:513–24.

41. Aedo S, Tomasz A. 2016. Role of the Stringent Stress Response in the Antibiotic Resistance Phenotype of Methicillin-Resistant Staphylococcus aureus. Antimicrob Agents Chemother 60:2311–7.

42. Geiger T, Goerke C, Fritz M, Schafer T, Ohlsen K, Liebeke M, Lalk M, Wolz C. 2010. Role of the (p)ppGpp synthase RSH, a RelA/SpoT homolog, in stringent response and virulence of Staphylococcus aureus. Infect Immun 78:1873–83.

43. Lemos JA, Lin VK, Nascimento MM, Abranches J, Burne RA. 2007. Three gene products govern (p)ppGpp production by Streptococcus mutans. Mol Microbiol 65:1568–81.

44. Dordel J, Kim C, Chung M, Pardos de la Gándara M, Holden MT, Parkhill J, de Lencastre H, Bentley SD, Tomasz A. 2014. Novel determinants of antibiotic resistance: identification of mutated loci in highly methicillin-resistant subpopulations of methicillin-resistant Staphylococcus aureus. mBio 5:e01000.

45. Martín-Rodríguez AJ, Römling U. 2017. Nucleotide Second Messenger Signaling as a Target for the Control of Bacterial Biofilm Formation. Curr Top Med Chem.

46. Liu A, Tran L, Becket E, Lee K, Chinn L, Park E, Tran K, Miller JH. 2010. Antibiotic sensitivity profiles determined with an Escherichia coli gene knockout collection: generating an antibiotic bar code. Antimicrobial agents and chemotherapy 54:1393–1403.

47. Blair JMA, Webber MA, Baylay AJ, Ogbolu DO, Piddock LJV. 2015. Molecular mechanisms of antibiotic resistance. Nature Reviews Microbiology 13:42–51.

48. Turner NA, Sharma-Kuinkel BK, Maskarinec SA, Eichenberger EM, Shah PP, Carugati M, Holland TL, Fowler VG. 2019. Methicillin-resistant Staphylococcus aureus: an overview of basic and clinical research. Nature Reviews Microbiology 17:203–218.

49. Worthington RJ, Melander C. 2013. Combination approaches to combat multidrug-resistant bacteria. Trends in biotechnology 31:177–184.

50. Waters EM, Rudkin JK, Coughlan S, Clair GC, Adkins JN, Gore S, Xia G, Black NS, Downing T, O’Neill E, Kadioglu A, O’Gara JP. 2017. Redeploying beta-lactam antibiotics as a novel antivirulence strategy for the treatment of methicillin-resistant Staphylococcus aureus infections. J Infect Dis 215:80–87.

51. Plata KB, Riosa S, Singh CR, Rosato RR, Rosato AE. 2013. Targeting of PBP1 by beta-lactams determines recA/SOS response activation in heterogeneous MRSA clinical strains. PLoS One 8:e61083.

52. Fey PD, Endres JL, Yajjala VK, Widhelm TJ, Boissy RJ, Bose JL, Bayles KW. 2013. A genetic resource for rapid and comprehensive phenotype screening of nonessential Staphylococcus aureus genes. mBio 4:e00537–12.

53. Chin D, Goncheva MI, Flannagan RS, Heinrichs DE. 2021. Mutations in a Membrane Permease or hpt Lead to 6-Thioguanine Resistance in Staphylococcus aureus. Antimicrob Agents Chemother 65:e0076021.

54. Salamaga B, Kong L, Pasquina-Lemonche L, Lafage L, von Und Zur Muhlen M, Gibson JF, Grybchuk D, Tooke AK, Panchal V, Culp EJ, Tatham E, O’Kane ME, Catley TE, Renshaw SA, Wright GD, Plevka P, Bullough PA, Han A, Hobbs JK, Foster SJ. 2021. Demonstration of the role of cell wall homeostasis in Staphylococcus aureus growth and the action of bactericidal antibiotics. Proc Natl Acad Sci U S A 118.

55. Aiba Y, Katayama Y, Hishinuma T, Murakami-Kuroda H, Cui L, Hiramatsu K. 2013. Mutation of RNA polymerase β-subunit gene promotes heterogeneous-to-homogeneous conversion of β-lactam resistance in methicillin-resistant Staphylococcus aureus. Antimicrob Agents Chemother 57:4861–71.

56. Gao W, Cameron DR, Davies JK, Kostoulias X, Stepnell J, Tuck KL, Yeaman MR, Peleg AY, Stinear TP, Howden BP. 2012. The RpoB H481Y Rifampicin Resistance Mutation and an Active Stringent Response Reduce Virulence and Increase Resistance to Innate Immune Responses in Staphylococcus aureus. The Journal of Infectious Diseases 207:929–939.

57. Gao W, Cameron DR, Davies JK, Kostoulias X, Stepnell J, Tuck KL, Yeaman MR, Peleg AY, Stinear TP, Howden BP. 2013. The RpoB H(4)(8)(1)Y rifampicin resistance mutation and an active stringent response reduce virulence and increase resistance to innate immune responses in Staphylococcus aureus. J Infect Dis 207:929–39.

58. Johnson S, Krüger D, Labischinski H. 1995. FemA of Staphylococcus aureus: isolation and immunodetection. FEMS Microbiol Lett 132:221–8.

59. Vilhelmsson O, Miller KJ. 2002. Synthesis of pyruvate dehydrogenase in Staphylococcus aureus is stimulated by osmotic stress. Appl Environ Microbiol 68:2353–8.

60. Li L, Cheung A, Bayer AS, Chen L, Abdelhady W, Kreiswirth BN, Yeaman MR, Xiong YQ. 2016. The Global Regulon sarA Regulates beta-Lactam Antibiotic Resistance in Methicillin-Resistant Staphylococcus aureus In Vitro and in Endovascular Infections. J Infect Dis 214:1421–1429.

61. Bowman L, Zeden MS, Schuster CF, Kaever V, Gründling A. 2016. New Insights into the Cyclic Di-adenosine Monophosphate (c-di-AMP) Degradation Pathway and the Requirement of the Cyclic Dinucleotide for Acid Stress Resistance in Staphylococcus aureus. J Biol Chem 291:26970–26986.

62. Boonsiri T, Watanabe S, Tan X-E, Thitiananpakorn K, Narimatsu R, Sasaki K, Takenouchi R, Sato’o Y, Aiba Y, Kiga K, Sasahara T, Taki Y, Li F-Y, Zhang Y, Azam AH, Kawaguchi T, Cui L. 2020. Identification and characterization of mutations responsible for the β-lactam resistance in oxacillin-susceptible mecA-positive Staphylococcus aureus. Scientific Reports 10:16907.

63. Sause WE, Balasubramanian D, Irnov I, Copin R, Sullivan MJ, Sommerfield A, Chan R, Dhabaria A, Askenazi M, Ueberheide B, Shopsin B, Bakel Hv, Torres VJ. 2019. The purine biosynthesis regulator PurR moonlights as a virulence regulator in <i>Staphylococcus aureus</i>. Proceedings of the National Academy of Sciences 116:13563–13572.

64. Goncheva MI, Flannagan RS, Sterling BE, Laakso HA, Friedrich NC, Kaiser JC, Watson DW, Wilson CH, Sheldon JR, McGavin MJ, Kiser PK, Heinrichs DE. 2019. Stress-induced inactivation of the Staphylococcus aureus purine biosynthesis repressor leads to hypervirulence. Nat Commun 10:775.

65. Gallagher LA, Shears R, Fingleton C, Alvarez L, M. WE, Clarke J, Bricio-Moreno L, Campbell C, Yadav A, Razvi F, O’Neill E, Cava F, O’Neill AJ, Fey PD, Kadioglu A, O’Gara JP. 2019. Impaired alanine transport or exposure to D-cycloserine increases the susceptibility of MRSA to beta-lactam antibiotics. bioRxiv 616920 doi: https://doi.org/10.1101/616920)..

66. Gélinas M, Museau L, Milot A, Beauregard PB. 2020. Cellular adaptation and the importance of the purine biosynthesis pathway during biofilm formation in Gram-positive pathogens. bioRxiv doi:10.1101/2020.12.11.422287:2020.12.11.422287.

67. Chin D, Goncheva MI, Flannagan RS, Deecker SR, Guariglia-Oropeza V, Ensminger AW, Heinrichs DE. 2021. Coagulase-negative staphylococci release a purine analog that inhibits Staphylococcus aureus virulence. Nat Commun 12:1887.

68. Baek KT, Jensen C, Farha MA, Nielsen TK, Paknejadi E, Mebus VH, Vestergaard M, Brown ED, Frees D. 2021. A Staphylococcus aureus clpX Mutant Used as a Unique Screening Tool to Identify Cell Wall Synthesis Inhibitors that Reverse beta-Lactam Resistance in MRSA. Front Mol Biosci 8:691569.

69. Giulieri SG, Guerillot R, Kwong JC, Monk IR, Hayes AS, Daniel D, Baines S, Sherry NL, Holmes NE, Ward P, Gao W, Seemann T, Stinear TP, Howden BP. 2020. Comprehensive Genomic Investigation of Adaptive Mutations Driving the Low-Level Oxacillin Resistance Phenotype in Staphylococcus aureus. mBio 11.

70. Fingleton C, Zeden MS, Bueno E, Cava F, O’Gara JP. 2021. Mutation of lipoprotein processing pathway gene <em>lspA</em> or inhibition of LspA activity by globomycin increases MRSA resistance to β-lactam antibiotics. bioRxiv doi:10.1101/2021.02.03.429649:2021.02.03.429649.

71. CLSI. 2018. Performance Standards for Antimicrobial Disk Susceptibility Tests; approved standard—12th ed. M02-A13., Wayne, PA.

72. Wayne P. CLSI 2018. CLSI 2018. Performance standards for antimicrobial disk susceptibility tests. Approved standard, 12th ed. CLSI standard M02-A13. Clinical and Laboratory Standards Institute,.

73. Underwood AJ, Zhang Y, Metzger DW, Bai G. 2014. Detection of cyclic di-AMP using a competitive ELISA with a unique pneumococcal cyclic di-AMP binding protein. J Microbiol Methods 107:58–62.

74. Saiman L, O’Keefe M, Graham PL, 3rd, Wu F, Said-Salim B, Kreiswirth B, LaSala A, Schlievert PM, Della-Latta P. 2003. Hospital transmission of community-acquired methicillin-resistant Staphylococcus aureus among postpartum women. Clin Infect Dis 37:1313–9.

75. O’Neill E, Pozzi C, Houston P, Smyth D, Humphreys H, Robinson DA, O’Gara JP. 2007. Association between methicillin susceptibility and biofilm regulation in Staphylococcus aureus isolates from device-related infections. J Clin Microbiol 45:1379–88.

76. Robinson DA, Enright MC. 2003. Evolutionary models of the emergence of methicillin-resistant Staphylococcus aureus. Antimicrob Agents Chemother 47:3926–34.

77. Conlon KM, Humphreys H, O’Gara JP. 2002. icaR encodes a transcriptional repressor involved in environmental regulation of ica operon expression and biofilm formation in Staphylococcus epidermidis. J Bacteriol 184:4400–8.

78. Gill SR, Fouts DE, Archer GL, Mongodin EF, Deboy RT, Ravel J, Paulsen IT, Kolonay JF, Brinkac L, Beanan M, Dodson RJ, Daugherty SC, Madupu R, Angiuoli SV, Durkin AS, Haft DH, Vamathevan J, Khouri H, Utterback T, Lee C, Dimitrov G, Jiang L, Qin H, Weidman J, Tran K, Kang K, Hance IR, Nelson KE, Fraser CM. 2005. Insights on evolution of virulence and resistance from the complete genome analysis of an early methicillin-resistant Staphylococcus aureus strain and a biofilm-producing methicillin-resistant Staphylococcus epidermidis strain. J Bacteriol 187:2426–38.

79. Horsburgh MJ, Aish JL, White IJ, Shaw L, Lithgow JK, Foster SJ. 2002. sigmaB modulates virulence determinant expression and stress resistance: characterization of a functional rsbU strain derived from Staphylococcus aureus 8325-4. J Bacteriol 184:5457–67.

