## Supplemental Figure 1 for "Purine nucleosides interfere with c-di-AMP levels and act as adjuvants to re-sensitise MRSA to β-lactam antibiotics"

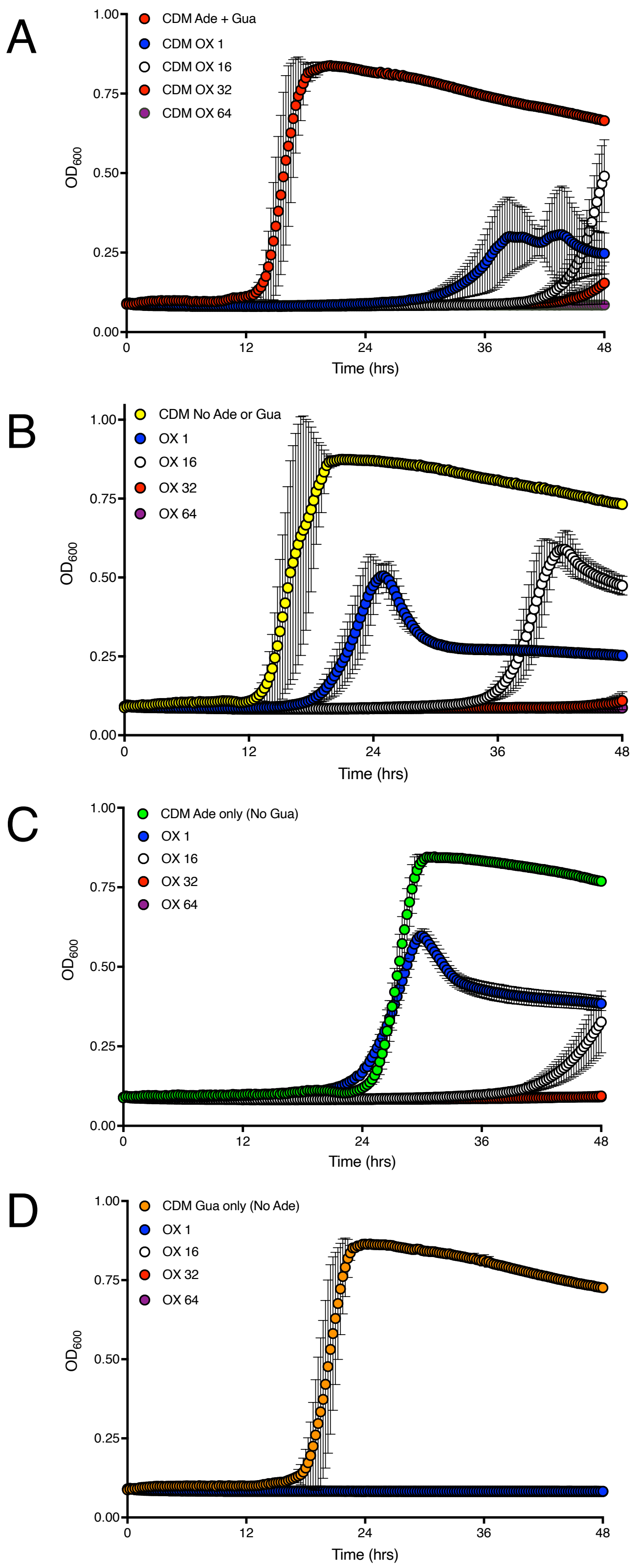


**Fig. S1. Exogenous guanine and adenine control oxacillin resistance of wild-type JE2 grown in modified chemically defined media. A.** Growth of JE2 for 48 hrs at 35ºC in modified chemically defined media without glucose (CDM) supplemented with 40 mg/l of both adenine and guanine (CDM Ade + Gua) and oxacillin (OX) 1, 16, 32 or 64 μg/ml. **B.** Growth of JE2 in CDM No Ade or Gua supplemented with OX 1, 16, 32 or 64 μg/ml. **C.** Growth of JE2 in CDM Ade only (no Gua) supplemented with OX 1, 16, 32 or 64 μg/ml. **D.** Growth of JE2 in CDM Gua only (no Ade) supplemented with OX 1, 16, 32 or 64 μg/ml. Growth ($OD_{600}$) was measured at 15 min intervals in a Tecan plate reader. Data are the average of 3 independent experiments and error bars represent standard deviation.
