## Supplemental Figure 2 for "Purine nucleosides interfere with c-di-AMP levels and act as adjuvants to re-sensitise MRSA to β-lactam antibiotics"

Oxacillin ($\mu$g/ml)

| Guanosine ($\mu$g/ml) | 0 | 0.0625 | 0.125 | 0.25 | 0.5 | 1 | 2 | 4 | 8 | 16 | 32 | 64 |
|---|---|---|---|---|---|---|---|---|---|---|---|---|
| 512 | 0.424 | 0.239 | 0.131 | 0.079 | 0.072 | 0.059 | 0.058 | 0.054 | 0.053 | 0.053 | 0.058 | 0.066 |
| 256 | 0.38 | 0.254 | 0.198 | 0.179 | 0.102 | 0.067 | 0.066 | 0.061 | 0.055 | 0.054 | 0.055 | 0.064 |
| 128 | 0.39 | 0.224 | 0.209 | 0.171 | 0.122 | 0.101 | 0.081 | 0.077 | 0.057 | 0.054 | 0.055 | 0.06 |
| 64 | 0.413 | 0.29 | 0.209 | 0.175 | 0.156 | 0.143 | 0.129 | 0.113 | 0.084 | 0.056 | 0.053 | 0.064 |
| 32 | 0.399 | 0.263 | 0.23 | 0.184 | 0.182 | 0.166 | 0.172 | 0.181 | 0.097 | 0.058 | 0.055 | 0.064 |
| 16 | 0.405 | 0.305 | 0.254 | 0.228 | 0.227 | 0.2 | 0.217 | 0.207 | 0.185 | 0.132 | 0.054 | 0.064 |
| 8 | 0.397 | 0.337 | 0.282 | 0.221 | 0.255 | 0.241 | 0.259 | 0.224 | 0.189 | 0.085 | 0.066 | 0.066 |
| 0 | 0.404 | 0.304 | 0.221 | 0.169 | 0.163 | 0.162 | 0.192 | 0.148 | 0.163 | 0.098 | 0.189 | 0.062 |

**Fig. S2. Guanosine increases the oxacillin susceptibility of MRSA strain JE2 in a concentration dependent manner.** Checkerboard titration assays were conducted using guanosine and oxacillin with JE2 grown for 24 h in Mueller Hinton 2% NaCl broth in 96-well plates. The data shown are the $A_{600}$ values for each well. The experiments were repeated at least three times and the data from a representative 96-well plate is shown. Purple shaded boxes indicated wells in which significant growth was measured.
