## Supplemental Figure 3 for "Purine nucleosides interfere with c-di-AMP levels and act as adjuvants to re-sensitise MRSA to β-lactam antibiotics"

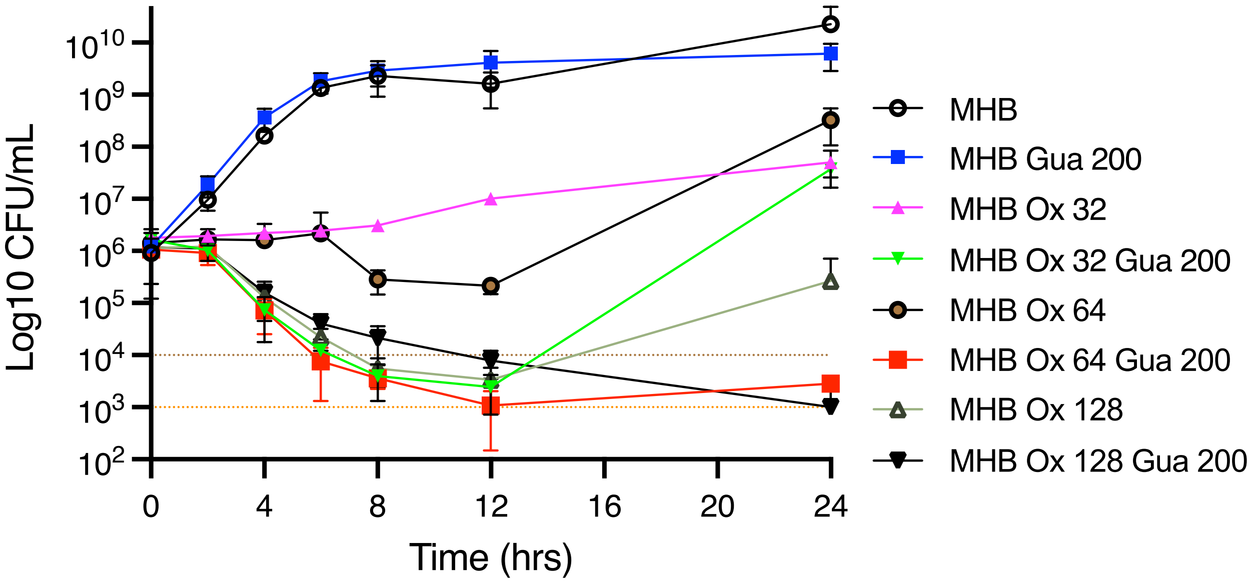


**Fig. S3.** *In vitro* kill curves for wild-type JE2 in MHB, MHB guanosine (Gua, 200 μg/ml), MHB oxacillin (OX, 32 μg/ml, 0.5× MIC; 64 μg/ml, 1.0× MIC; 128 μg/ml, 2.0× MIC) and combinations of Gua and OX. Three hour cultures were adjusted to $10^6$ CFU/ml in Mueller Hinton 2% NaCl broth (MHB, $A_{600}$=0.001) or MHB supplemented with Gua and/or OX (Ox). Cultures were incubated at 35ºC and the number of CFU/ml enumerated at 0, 2, 4, 6, 8, 12 and 24 h. The data presented are the mean of three independent experiments, and standard error of the mean is shown. Antibiotic synergism was defined as a ≥2 $\log^{10}$ decrease in the number of CFU/ml in JE2 cell suspensions exposed to OX/Gua combinations compared to Ox alone. Reductions in the number of CFU/ml to ≤$10^3$ was defined as bactericidal activity.
