## Supplemental Figure 4 for "Purine nucleosides interfere with c-di-AMP levels and act as adjuvants to re-sensitise MRSA to β-lactam antibiotics"

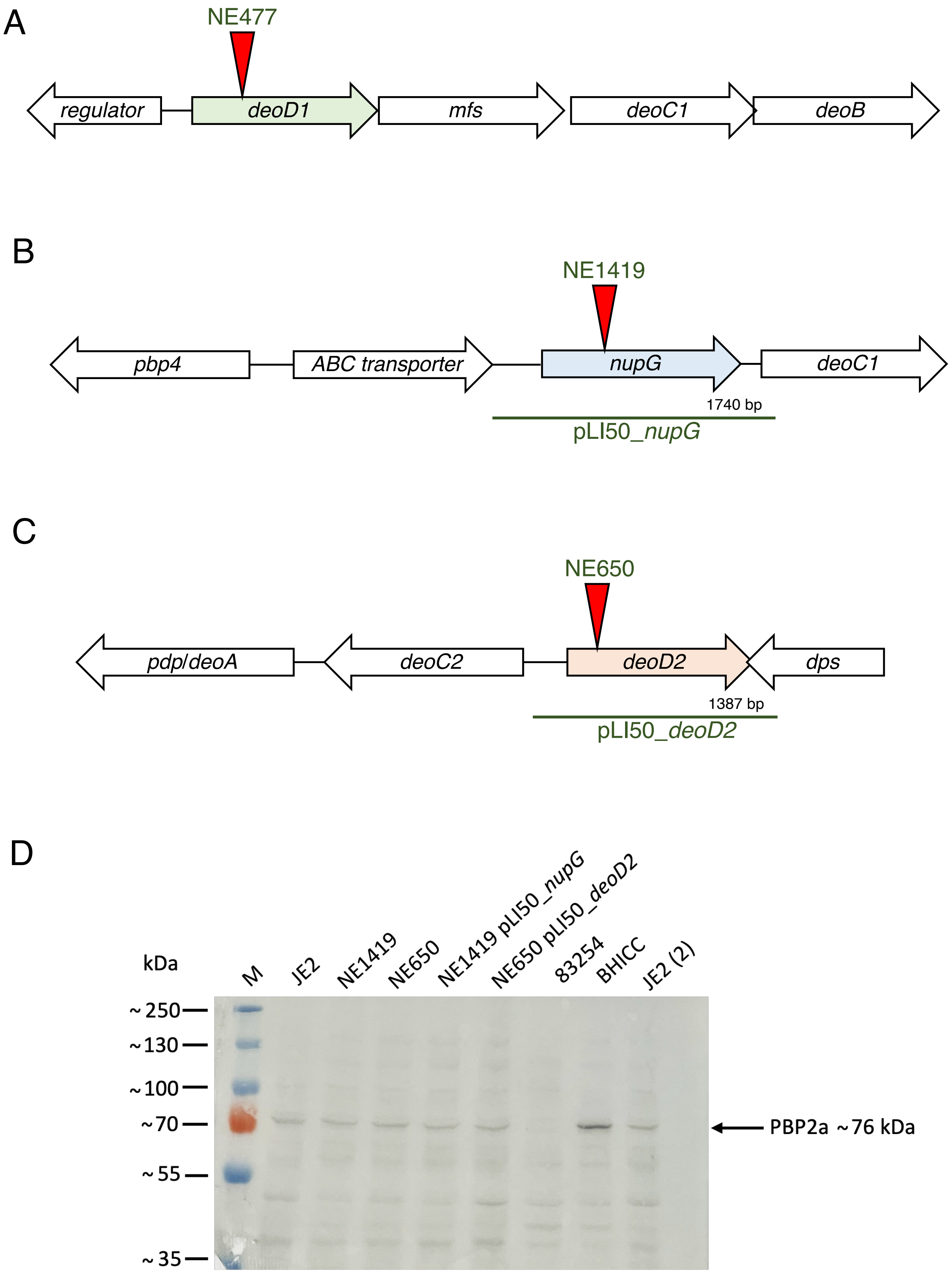

A
NE477
regulator
deoD1
mfs
deoC1
deoB
B
NE1419
pbp4
ABC transporter
nupG
deoC1
1740 bp
pLI50_nupG
C
NE650
pdp/deoA
deoC2
deoD2
dps
1387 bp
pLI50_deoD2
D
kDa
M
JE2
NE1419
NE650
NE1419 pLI50_nupG
NE650 pLI50_deoD2
83254
BHICC
JE2 (2)
~250
~130
~100
~70
~55
~35
PBP2a ~76 kDa

**Fig. S4. Chromosomal organization of *deoD1*, *nupG, and deoD2* and Western blot analysis of PBP2a in the NE1419 (*nupG*) and NE650 (*deoD2*) mutants. A.** Location of the transposon insertion in NE1419 (*nupG*::$Em^r$) and the chromosomal locus amplified and cloned into pLI50_*nupG* for complementation experiments. **D.** Location of the transposon insertion in NE650 (*deoD2*::$Em^r$) and the chromosomal locus amplified and cloned into pLI50_*deoD2* for complementation experiments. **C.** Location of the transposon insertion in NE477 (*deoD1*::$Em^r$). **D.** Western blot of PBP2a protein in wild-type JE2, NE1419 (*nupG*), NE650 (*deoD2*), NE1419 pLI50_*nupG*, NE650 pLI50_*deoD2*, MSSA strain 8325-4 (negative control), HoR MRSA strain BH1CC (positive control) and JE2 ($2^{nd}$ well). The strains were grown overnight in MHB 2% NaCl supplemented with 5 *µ*g/ml oxacillin (OX), except for BHICC which was grown in MHB NaCl supplemented with 32 µg/ml OX and 8325-4 which was grown in MHB NaCl with no OX. For each sample, 6 *µ*g total protein was run on a 7.5% Tris-Glycine gel, transferred to a PVDF membrane and probed with anti-PBP2a (1:1000), followed by HRP-conjugated protein G (1:2000) and colorimetric detection with Opti-4CN Substrate kit. Three independent experiments were performed, and a representative image is shown.
