## Supplemental Figure 5 for "Purine nucleosides interfere with c-di-AMP levels and act as adjuvants to re-sensitise MRSA to β-lactam antibiotics"

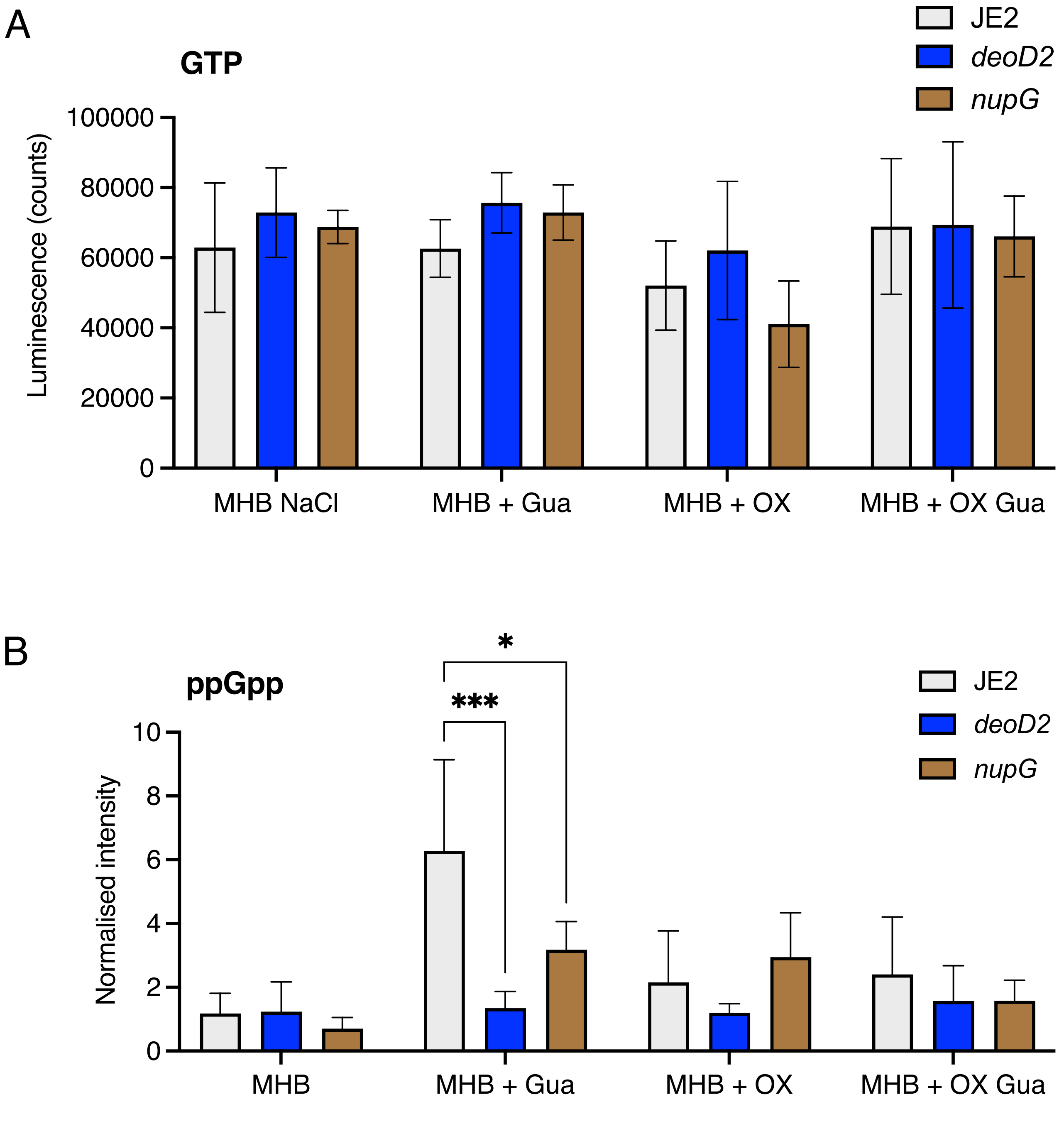


**Fig. S5. Impact of exogenous guanosine or mutations in *nupG* and *deoD* on GTP and ppGpp levels. A.** GTPase-Glo bioluminescence assay of GTP levels in JE2, *nupG* and *deoD2* grown in MHB NaCl supplemented with guanosine (Gua, 0.2 g/l) and/or oxacillin (OX, 1 μg/ml). **B.** Radiochemical thin layer chromatographic analysis of ppGpp levels in JE2, *nupG* and *deoD2* grown in MHB NaCl with Gua 0.2 g/l and/or OX 1 μg/ml. Data are the average of 3 independent experiments and error bars indicate standard deviation. Asterisks indicate statistically significant differences according to one-way ANOVA followed by Tukey's multiple comparison post-hoc test (* $p<0.05$, *** $p<0.001$).
