## Supplemental Figure 6 for "Purine nucleosides interfere with c-di-AMP levels and act as adjuvants to re-sensitise MRSA to β-lactam antibiotics"

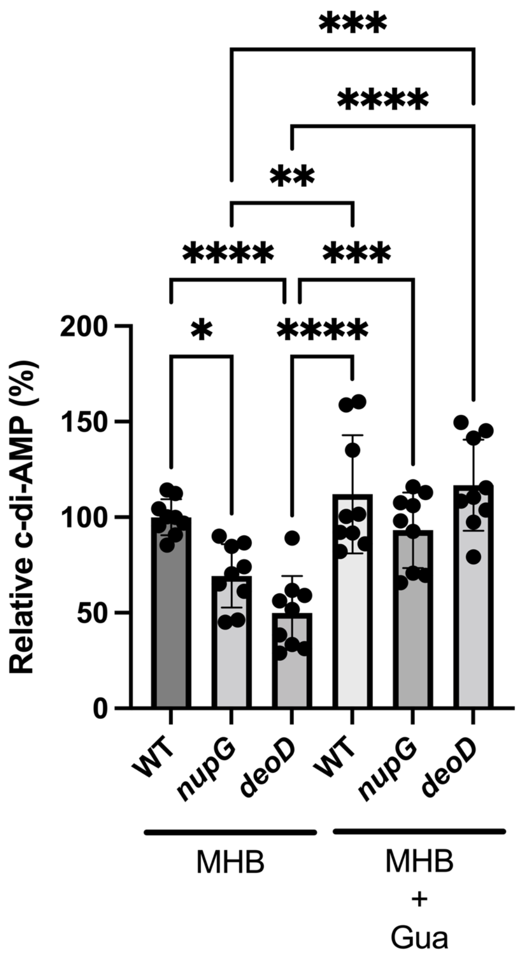


**Fig. S6.** Relative c-di-AMP levels in wild-type, *nupG* and *deoD2* grown for 18 h in 5 ml MHB cultures without 2% NaCl, with and without 0.2 g/l guanosine (Gua). c-di-AMP levels are the averages of nine biological replicates from a combination of three separate experiments with three bio-replicates each, normalized as % relative to wild type. Asterisks indicate statistically significant difference according to one-way ANOVA followed by Tukey's multiple comparison post-hoc test (* $p<0.05$, ** $p<0.01$, *** $p<0.001$, **** $p<0.0001$). Error bars indicate standard deviation.
